# Drought and Herbivory Shape Morphological and Chemical Traits in Black Poplar (*Populus nigra*)

**DOI:** 10.64898/2026.06.30.735551

**Authors:** Sarah K. Weirauch, Hanna Greßmann, Michael Reichelt, Elisabeth Kaltenegger, Jörg-Peter Schnitzler, Sybille B. Unsicker

## Abstract

Due to climate change, extreme weather events such as droughts are becoming more frequent and intense. This has a profound impact on plant performance and ecological interactions, including those involving herbivorous insects. The combined impact of drought stress and insect herbivory on plant metabolism has rarely been studied, particularly in woody plants. In this study, we investigated the influence of varying degrees of drought, both alone and in combination with herbivory by the leaf beetle *Chrysomela tremulae*, on the morphological and chemical characteristics of black poplar (*Populus nigra*) trees using a full factorial experimental design. We quantified morphological traits, volatile organic compound (VOC) emissions, phytohormone and amino acid concentrations, and phenolic profiles. Drought conditions increased the concentrations of salicylic acid (SA) and abscisic acid (ABA), while feeding induced ABA and SA. Amino acid profiles shifted significantly under drought conditions, particularly in beetle-infested plants. In contrast, salicinoids, which are the most important phenolic defense compounds in poplars, remained relatively stable. We also observed significant compound-specific effects on both constitutive and herbivore-induced VOC emissions. Our results demonstrate that drought and insect herbivory exert a joint influence on the chemical responses of *P. nigra* across multiple metabolic pathways. These findings highlight how the interaction between abiotic and biotic stresses can influence the defense chemistry of trees, which will consequently affect ecological interactions in forest ecosystems in the face of climate change.

## 1. Introduction

Climate change poses a major challenge to plants in both natural and managed ecosystems worldwide. More frequently occurring extreme weather conditions such as drought, heat, and flooding, together with rising atmospheric CO₂ concentrations, contribute to habitat fragmentation, altered plant resource availability, and destabilization of ecological communities (Gamfeldt et al., 2013; Rosenzweig et al., 2008; Weiskopf et al., 2020). When examining the consequences of climate change, forests are of particular concern because of their global economic and ecological importance. They store large amounts of biomass, act as essential carbon sinks, and provide important ecosystem services such as climate regulation, soil stabilization, nutrient cycling, and habitat provisioning (Brockerhoff et al., 2017; Dixon et al., 1994). In particular, drought as an abiotic stress is considered as one of the most influential drivers of forest ecosystem decline (Allen and Breshears, 1998; Bennett et al., 2015). Drought triggers a suite of coordinated plant responses, including stomatal regulation, osmotic adjustment, changes in biomass allocation, and shifts in secondary metabolism (Fàbregas and Fernie, 2019; Haghpanah et al., 2024). These responses include reduction of leaf area and overall growth as well as changes in stomatal conductance and root-to-shoot ratios depending on the drought intensity (Bansal et al., 2013). The threshold at which dry conditions elicit physiological drought stress varies among taxa, reflecting differences in adaptation (Kozlowski and Pallardy, 2002). Many temperate tree species are isohydric and maintain leaf water potential by rapidly closing stomata under water deficit, thereby limiting carbon assimilation (McDowell et al., 2008; Tardieu and Davies, 1992; Tardieu and Simonneau, 1998). In contrast, anisohydric plants regulate stomata more loosely, maintaining photosynthesis while allowing stronger fluctuations in leaf water potential (McDowell et al., 2008; Tardieu and Simonneau, 1998). Structural traits such as deeper root systems or altered leaf morphology further contribute to drought tolerance (Regier et al., 2009). Drought effects also differ strongly across developmental stages: younger trees often experience more severe growth reduction but show a higher capacity for recovery after rewatering than mature individuals (Au et al., 2022). Under natural conditions, trees rarely face one single stressor in isolation but experience a suite of stresses such as drought and insect herbivory at the same time. The interaction between these stressors is complex and may be additive, antagonistic, or synergistic depending on the intensity of drought, the herbivore involved, and the trait under investigation. Earlier studies have shown that drought can alter susceptibility to herbivores, modify compensatory growth responses, or intensify or dampen herbivore-induced chemical defenses (Bansal et al., 2013; Copolovici et al., 2014; Malone et al., 2023). Drought alters both the quantity and quality of metabolites involved in plant resistance, including signalling pathways associated with jasmonic and salicylic acid (JA and SA) as well as phenolic compounds (Ahmad et al., 2023). Trees have a range of direct and indirect defenses to cope with herbivory (Büchel et al., 2016; Gols, 2014; Gong and Zhang, 2014). Direct chemical tree defenses include phenolics like tannins, salicinoids, and other compounds that deter feeding or reduce herbivore performance. Indirect defenses include the emission of volatile organic compounds (VOCs), which serve important ecological functions like stress signalling, deterring insect herbivores, or attracting natural enemies of herbivores like parasitoids and predators (Heil, 2008). Tree VOCs encompass several biosynthetic classes, including terpenoids, green leaf volatiles (GLVs), benzenoids, and nitrogen-containing compounds (Laothawornkitkul et al., 2009; Pichersky and Gershenzon, 2002). Here, monoterpenes and sesquiterpenes are the most abundant VOCs that play major roles in stress signalling and plant defense (Clavijo McCormick et al., 2014b; Dudareva et al., 2006, Unsicker et al. 2009). VOCs are metabolically costly and tightly regulated by resource availability. Environmental stress, especially drought, can substantially alter VOC emissions in a compound-specific manner (Peñuelas and Staudt, 2010; Reinecke et al., 2024). Moderate drought was found to increase emissions of stress-related volatiles such as GLVs, nitrogenous compounds, and certain terpenoids, whereas severe or prolonged drought typically suppresses emissions by limiting *de novo* synthesis (Brilli et al., 2007). In addition to directly affecting metabolic processes, drought-induced changes in plant morphology and leaf structure may further influence herbivore feeding behavior and patterns of volatile release, adding additional layers of complexity. Moreover, trees differ fundamentally from many model plants due to their long lifespan, large biomass, and investment in constitutive and inducible chemical defenses (Coley et al., 1985; Haukioja and Koricheva, 2000). Consequently, drought and herbivory may interact differently in woody species compared with herbaceous plants. Responses to abiotic and biotic stress can also vary among genotypes, and genetic background has been shown to contribute strongly to phytochemical diversity and stress responsiveness (Eisenring et al., 2021; Haviola et al., 2012; Thakur et al., 2021). However, studies that simultaneously address morphological responses, defense metabolites, and VOC emissions under combined drought and herbivory in trees remain comparatively scarce.

The aim of this study was to investigate how different drought intensities and herbivory by the specialist leaf beetle *Chrysomela tremulae* shape morphological traits, leaf and root chemistry, and VOC emissions, in young *Populus nigra* (black poplar) trees. We used *P. nigra* as our model tree species. As a fast-growing riparian tree adapted to both flooding- and drought-prone habitats, it experiences fluctuating water availability in nature. Poplar trees emit high levels of isoprene, and herbivory has been shown to strongly increase the emission of monoterpenes and sesquiterpenes (Clavijo McCormick et al., 2014a; Sasaki et al., 2005). Nitrogen-containing compounds, aromatics, and GLVs contribute substantially to signalling under biotic and abiotic stress in poplar (Clavijo McCormick et al., 2019). Moreover, salicinoids such as salicin, salicortin, and their derivatives play a central role in chemical defense in *Populus* and respond to water stress and herbivory (Boeckler et al., 2013; Hale et al., 2005). Using young *P. nigra* trees of three different genotypes and herbivory by the specialist leaf beetle *Chrysomela tremulae,* we addressed the following questions: (1) Does water stress (drought) act as a primary driver of plant phenotypes, inducing pronounced changes in plant morphology and metabolite profiles? (2) Is herbivore-induced VOC emission shaped by drought intensity, such that the magnitude and composition of volatile responses depend on plant water status? (3) Do drought and herbivory interact in shaping morphological and chemical traits, resulting in synergistic or antagonistic responses rather than additive effects? Together, these questions allow us to assess how water stress and herbivory influence plant morphology, defense-related metabolites, and VOC emissions in *P. nigra*.

## 2. Materials and Methods

### 2.1 Plants and Insects

Three different genotypes of *Populus nigra* (black poplar, Salicaceae), obtained from a tree nursery (Waller Baumschulen, Schwäbisch Hall, Germany), were grown from stem cuttings in a 1:1 mixture of sand and soil (Klasmann potting substrate; Klasman-Deilmann, Geeste, Germany). They were cultivated in 2 L pots under summer conditions (21 °C, 16 h/8 h light cycle) in a greenhouse, well-watered and fertilized once per week with a 0.1 % Ferty 3 solution (Planta Düngemittel GmbH, 93123 Regenstauf). At the start of the experiment the plants were five months old.

*Chrysomela tremulae* (small poplar leaf beetle, *Chysomelidae*) beetles were obtained from our lab breeding at Kiel University. They were fed on leaves from young *P. nigra* trees. For this experiment, we used adult, egg-laying beetles with a balanced sex ratio.

### 2.2 Experimental setup and stress induction

Trees were assigned to six treatment groups in a full factorial design with three levels of water availability and two levels of experimental leaf damage (herbivory and control) (**Fig. S1**). Just before the onset of insect herbivory, trees were arranged in six blocks, each consisting of one plant per genotype and treatment combination (i.e., 18 plants per block), resulting in six replicates per genotype × treatment combination. All plants were placed in individual coasters to allow precise watering of the respective treatments. Water stress levels were defined relative to field capacity and similar to previous studies in woody plants categorized into three intensity classes at 100 % (approximately 50-60 % volumetric soil moisture), 50 % (15-20 %) and < = 25 % (< 15 %) field capacity (FC) respectively (Chen et al., 2010; Cocozza et al., 2010; Yin et al., 2005); (**Table S1**). Field capacity of the substrate was determined experimentally prior to the start of the experiment and served as a reference for all subsequent watering regimes, with volumetric soil water content monitored using a ML3 ThetaProbe sensor (Delta-T Devices, Cambridge, UK) calibrated for organic soil. Throughout the experimental phase, pots were weighed daily to track water loss, and water was added as required to maintain target moisture levels. Herbivory was induced by exposing trees to five adults of *C. tremulae* for two days. For this purpose, the first five fully expanded leaves per plant were enclosed in fine-mesh nets two days prior to volatile collection. Five individuals of *C. tremulae* were released into these nets on all trees in the herbivory treatment groups. Insects remained on the plants throughout the exposure period to ensure consistent and treatment-specific feeding.

### 2.3 Volatile collection and analysis

VOC collection from leaves of *P. nigra* trees started 10 days after the initiation of water stress treatments. Immediately before volatile sampling, beetles were removed from the nets. The same five leaves used for herbivory treatments were then enclosed in PET bags (Toppits® Bratschlauch, Melitta, Minden, Germany, **⌀** =30 cm). The bags were secured to the trunk using resealable cable ties. VOCs were then collected for 2 hrs using a mobile push-pull system as described earlier by Medina-van Berkum et al. (2026) with slight modifications. Briefly, charcoal-purified air was pumped into the bags at a flow rate of 1 L min⁻¹ and sucked out with another pump at 0.5 L min⁻¹ by passing a volatile collection trap packed with PORAPAK-Q™ (Analytical Research Systems, Inc., Gainesville, FL, USA). Volatile measures were performed in the experimental blocks described above. Volatile collection traps were eluted with 200 µl dichloromethane containing 10 ng μL⁻¹ nonyl acetate as an internal standard (SigmaAldrich, St. Louis, MO, USA). VOC measurements with GC-MS/FID were performed as described earlier (Clavijo Mccormick et al., 2014b; Eberl et al., 2018). Briefly, VOCs were analyzed with a gas chromatograph (Agilent 6890 series, Agilent, Santa Clara, CA, USA) coupled to either an Agilent 5973 series mass spectrometer (GC-MS, MS, temperatures: 270 °C (transfer line), 150 °C (quadrupole), 230 °C (source temperature); electron energy, 70 eV) or a flame ionization detector (FID) running at 300 °C. A DB-5MS column (30 m x 0.25 mm x 0.25 μm) with either He (for MS) or H_2_ (for FID) as carrier gases. The injection of the samples (1 µL; splitless) was carried out with a flow of 2 ml min^-1^ (oven starting temperature: 45 °C, increase to 180 °C at 6 °C min^-1^ and then to final temperature at 300 °C at 100 °C min^-1^). Compounds were identified by comparing their mass spectra data with mass spectral data from the NIST database. For the quantification of the compounds, the FID peak areas were calculated in relation to the peak areas of the internal standard. A detailed description of the comparison of measured and reference spectra including Match Number and Matching Probability can be found in **Table S9**.

### 2.4 Determining morphological traits and biomass harvest

To determine fresh weight, dry weight and leaf water content (calculated as (fresh weight - dry weight) / fresh weight × 100), the leaves from which the VOCs were collected were cut and weighed without their petioles. Afterwards, they were spread on a whiteboard with a 1 × 1 cm reference square and photographed. Leaf area and herbivore-inflicted leaf area loss was quantified as described in Boeckler et al. (2013) using Adobe Photoshop CS2 (Adobe Systems, San Jose, CA, USA). Thereafter, the leaves were halved with scissors and the midribs were removed. The halves of the leaves were placed alternately in one of two 5 ml vials, flash-frozen in liquid nitrogen and stored at -80 °C until further analysis. During harvesting, the roots were carefully removed from the substrate and washed with tap water until all adhering soil was completely removed. After quickly drying them with paper towels, roots were weighed, transferred to 50 mL tubes, flash-frozen in liquid nitrogen and then also stored at -80 °C. For dry weight determination and further processing, leaf and root samples were lyophilized for 48 hours (Alpha 1-4 LSC basic, Christ GmbH, Osterode, Germany). Leaf number and tree height were measured before the start of treatment and right before harvesting. The leaves that were not inside the bag for volatile collection were weighed and put in an oven at 70 °C for at least 48 hours before the dry weight was measured.

### 2.5 Chemical analyses of phenolic compounds

Lyophilized leaf and root material was homogenised using a paint shaker Merris Minimix (Merris, Ichtershausen, Germany). Phenolic compounds were extracted with 1 mL 100 % MeOH from 10 mg of finely ground, lyophilized leaf and root material. Phenyl-β-glucopyranoside (0.8 mg mL⁻¹) (Sigma-Aldrich, Deisenhofen, Germany) was added to each sample as an internal standard. After vortexing and incubation on a horizontal shaker for 30 min, samples were centrifuged (3 min, 3200 rpm). For HPLC analysis, 200 µL of the raw extract was diluted with 200 µl Milli-Q H₂O.

Phenolic analytes were separated and quantified using high-performance liquid chromatography with ultraviolet detection (HPLC-UV; Agilent Infinity III, Agilent Technologies, Santa Clara, CA, USA). Detection was carried out at 200 nm using a photodiode array (PDA) detector. Chromatographic separation was performed on an EC 250 × 4.6 mm NUCLEODUR Sphinx RP column (5 µm; Macherey-Nagel, Germany) connected to a C18 pre-column (5 µm, 4 × 3 mm; Phenomenex, USA). The injection volume was 20 µL, and the column temperature was maintained at 25 °C with a constant flow rate of 1 mL min⁻¹. The mobile phase consisted of solvent A (Milli-Q H₂O) and solvent B (acetonitrile), applied in gradient mode as follows (time [min] / % B): 0.00/14, 22.00/58, 22.10/100, 25.00/100, 25.10/14, 30.00/14. Phenolic compounds identified with commercially available authentic standards for salicin and catechin (Sigma-Aldrich, Merck KGaA, Darmstadt, Germany) and standards isolated and identified in an earlier study (Boeckler et al., 2013) are reported as relative amounts normalized to sample dry mass.

### 2.6 Amino acids and phytohormone analysis

Free amino acids and phytohormones were quantified in methanol extracts with a LC-MS/MS (Agilent 1260 Infinity II HPLC coupled to a QTRAP® 6500+ triple quadrupole mass spectrometer; Agilent Technologies, Santa Clara, CA, USA; AB Sciex, Waltham, MA, USA). Compounds were separated on a ZORBAX Eclipse XDB-C18 column (1.8 µm, 4.6 × 50 mm; Agilent Technologies) in multiple reaction monitoring (MRM) mode, with compound-specific methods and internal standards as described in Yepes-Vivas et al. (2025) and Alipour et al., (2025). For free amino acid analysis, methanolic extracts were diluted with water containing an isotopically labeled internal standard mixture (U13C, U15N-labeled algal amino acid mix, 10 µg mL⁻¹; Isotec, Miamisburg, OH, USA) and D5-tryptophan (5 µg mL⁻¹; Cambridge Isotope Laboratories, Andover, MA, USA). The mobile phases included solvent A (0.05 % formic acid in water (v:v)) and solvent B (acetonitrile). The chromatographic gradient was set as follows (time in min/concentration of solvent B in %): 0.00/3, 1.00/3, 2.70/100, 3.00/100, 3.10/3, 6.00/3. The constant flow rate was 1100 µL min^-1^, and the column oven temperature was maintained at 25 °C. Ionization occurred in positive electrospray ionization mode. Individual amino acids were quantified against their respective isotopically labeled standards.

Phytohormones were quantified from the methanolic raw extracts using compound-specific internal standards, including D₄-salicylic acid (Santa Cruz Biotechnology, Dallas, TX, USA), D₆-jasmonic acid (HPC Standards, Cunnersdorf, Germany), D₆-abscisic acid (Toronto Research Chemicals, Toronto, Canada), and D₆-jasmonoyl-isoleucine (HPC Standards, Germany). The mobile phases included solvent A (0.05 % formic acid in water (v:v)) and solvent B (acetonitrile). The chromatographic gradient was set as follows (time in min/concentration of solvent B in %): 0.00/10, 0.5/10, 4.0/90, 4.02/100, 4.5/100, 4.51/10, 7.00/10. The constant flow rate was 1100 µL min^-1^, and the column oven temperature was maintained at 25 °C. Ionization occurred in negative electrospray ionization mode. Jasmonates are reported as the sum of JA, JA-Ile, OH-JA, OH-JA-Ile, and COOH-JA-Ile. Detailed information on MRM transitions is provided in the supporting information (Phytohormones, **Table S2**; Amino acids, **Table S3**).

### 2.7 Statistical analysis

Statistical analyses were performed in R v4.4.3 (R Core Team, 2024). Model assumptions like normal distribution and homogeneity of variances were checked. Whenever necessary, response variables were log- or square-root transformed. Morphological traits and metabolite concentrations were analysed using linear mixed-effects models (LMMs). Water regime, herbivory, and their interaction were specified as fixed effects, and genotype was included as a random factor. The inclusion of block as a random effect was evaluated and removed when variance components were negligible or resulted in singular fits. For phenolic metabolites, biomass dry weight was included as a covariate to account for variation in plant growth and tissue investment. Treatment effects were assessed using Type III ANOVA with Satterthwaite approximation of degrees of freedom. For single VOC analyses, p-values were adjusted for false discovery rate (FDR) using the Benjamini-Hochberg procedure. Where significant effects were detected, Tukey-adjusted pairwise comparisons based on estimated marginal means were conducted. Relationships between herbivore damage intensity and VOC emissions were analysed within the mixed-effects modelling framework using herbivory-treated plants only. Statistical significance was assumed at p < 0.05. Model explanatory power was quantified using marginal and conditional R² following (Nakagawa and Schielzeth, 2013). Functional chemical diversity of VOC blends was quantified using Functional Hill diversity (q = 1), which integrates compound abundance and chemical dissimilarity among VOCs. In contrast to conventional Hill diversity metrics based solely on compound abundance, this approach incorporates functional trait information derived from chemical structure (SMILES and InChIKey identifiers) following (Petrén et al., 2023). Multivariate variation in free amino acid profiles was visualized using a heatmap of log10-transformed relative changes compared with the constitutive well-watered control. Data processing, statistical analyses, and visualisation employed the packages *lme4* (Bates et al., 2015), *lmerTest* (Kuznetsova et al., 2017), *emmeans* (Lenth, 2024), *performance* (Lüdecke et al., 2021), *DHARMa* (Hartig, 2022), *chemodiv* (Petrén et al., 2023), *pheatmap* (Kolde, 2019), *tidyverse* (Wickham et al., 2019) and *ggplot2* (Wickham, 2011). Final figure adjustments were performed using Inkscape v1.4.2. (Inkscape Project, 2025).

## 3. Results

### 3.1 Water stress reduces growth and alters biomass accumulation in *P. nigra*

Growth and biomass accumulation in *P. nigra* was significantly decreased in moderate and severe drought stress. Trees were highest in the well-watered situation and strongly reduced under both drought treatments, with no significant difference between moderate and severe drought levels (F = 33.14, p < 0.001; **Fig. 1**).

**Fig. 1:**
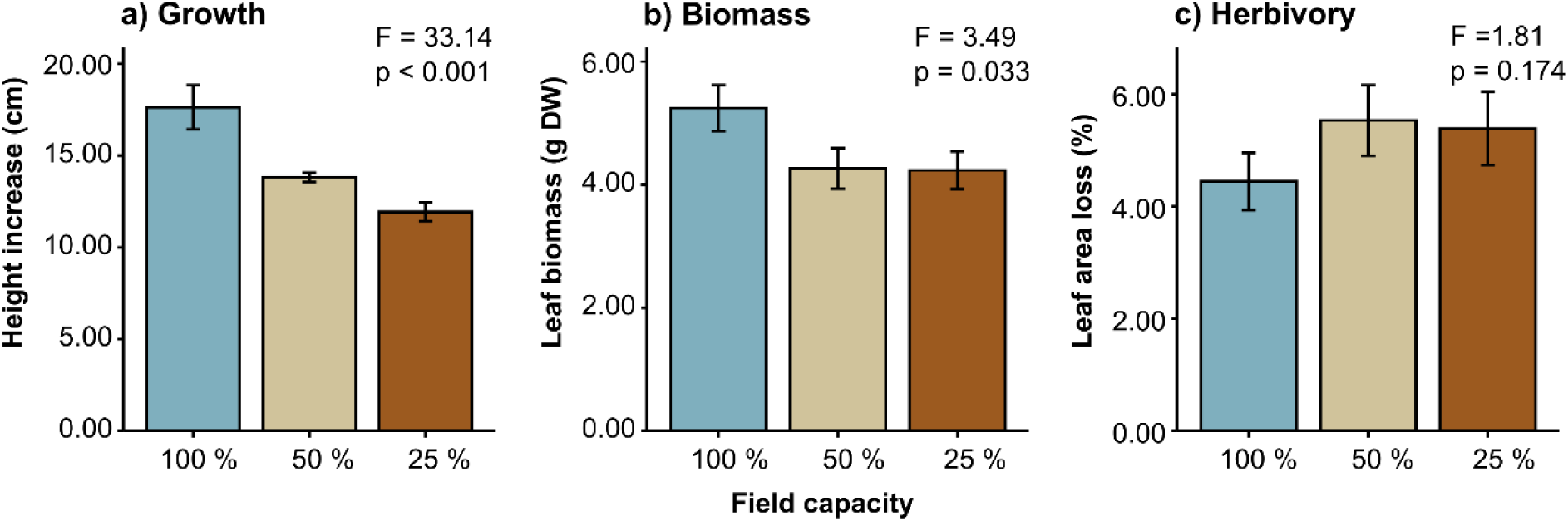
Effect of drought stress on morphological traits and experimental leaf beetle herbivory in young *Populus nigra* trees. (a) Height increase, (b) leaf biomass (dry weight), and (c) leaf area loss (%) in herbivory-treated plants only. Colours indicate water regime (field capacity: 100 %, blue; 50 %, beige; 25 % brown). Height increase and leaf biomass (dry weight) were analysed across all plants and separated by drought level, whereas leaf area loss due to specialist *Chrysomela tremulae* beetle feeding for 44 hours was analysed exclusively for plants exposed to herbivory. Linear mixed-effects models tested the effects of water regime, herbivory, and their interaction with genotype as a random effect (Table S4). Bars show means ± SE (n = 16-18).

Leaf biomass also declined in drought stressed trees (F = 3.49, p = 0.033; **Fig. 1b**). Both moderate and severe drought resulted in significantly lower dry weight, without differences between the two drought levels. In herbivory-treated plants, leaf area loss showed no significant changes between control and drought stressed trees (F = 1.81, p = 0.174; **Fig. 1c**).

### 3.2 Phenolic metabolites in *P. nigra* show compound- and tissue-specific responses

Responses to water regime and herbivory differed among phenolic compounds and between leaves and roots (Table S4; **Fig. 2**). Figure 2 presents treatment effects on phenolic metabolites in leaves and roots. In leaves, herbivory significantly increased total salicinoid concentrations (F = 6.30, p = 0.014; **Fig. 2a**), whereas the water regime and the interaction were not significant. Salicortin concentrations did not differ among treatments. In contrast, water regime had a significant effect of 6′-O-benzoylsalicortin in leaves, resulting in slightly higher concentrations in the water-stressed trees (F = 7.05, p = 0.001; **Fig. 2c**), herbivory (F = 4.95, p = 0.028), and their interaction (F = 4.36, p = 0.015). Catechin concentrations increased strongly under drought conditions (F = 17.66, p < 0.001; **Fig. 2d**), while herbivory and the interaction were not significant. Rutin concentrations showed significant effects of both water regimes (F = 3.21, p = 0.044; **Fig. 2e**) and herbivory (F = 5.38, p = 0.022). In roots, phenolic responses were primarily driven by water regime. Total salicinoids (F = 4.27, p = 0.017; **Fig. 2a**), salicortin (F = 4.96, p = 0.009; **Fig. 2b**), and catechin (F = 5.15, p = 0.007; Fig. **2d**) concentrations were significantly lower under drought. In the roots, neither herbivory nor water regime × herbivory interactions were significant for any phenolic compound. Salicinoids showed comparatively minor variation among treatments, whereas catechin exhibited pronounced drought-related responses.

**Fig. 2:**
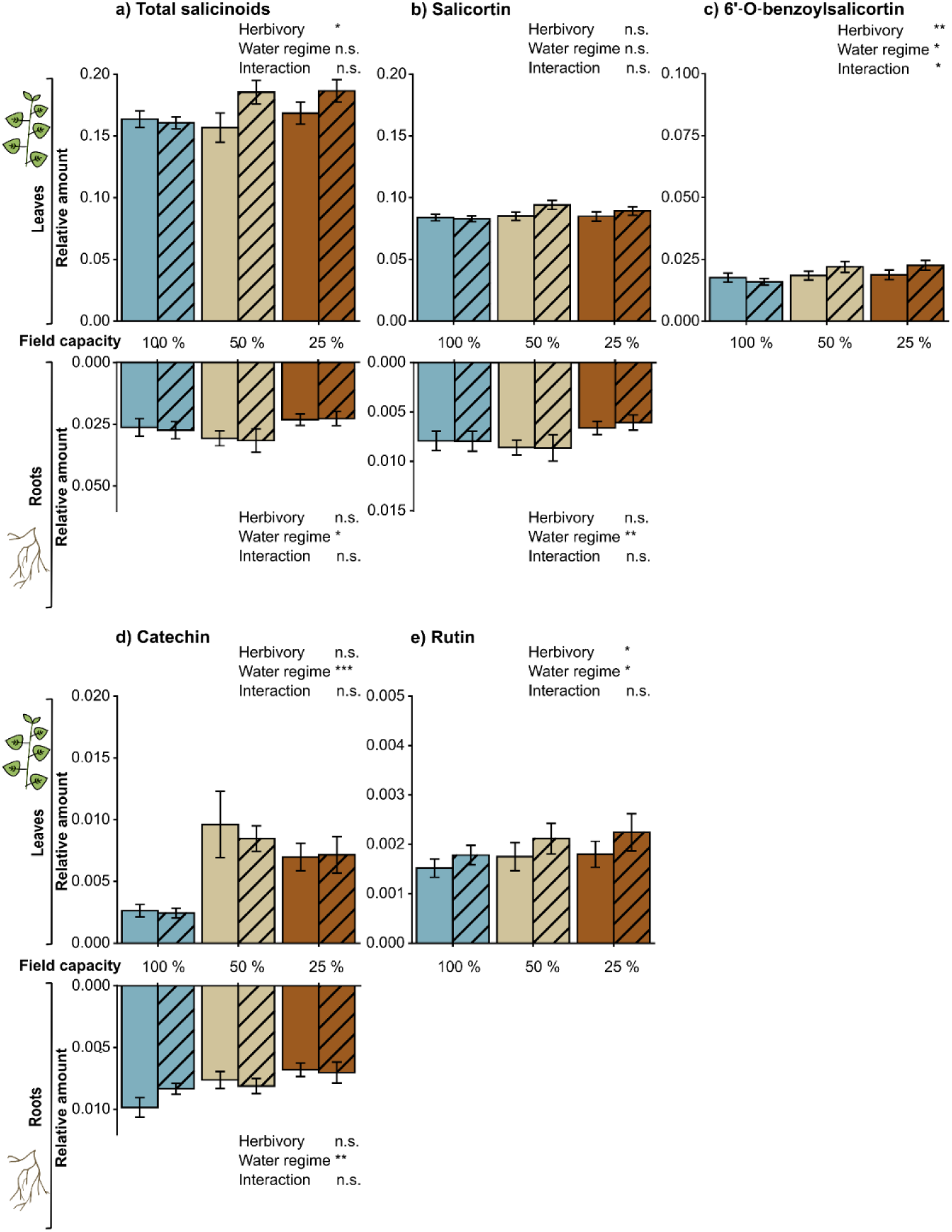
Effects of drought stress and herbivory on leaf and root phenolics in young *Populus nigra* trees. Salicinoids (a-c): (a) total salicinoids (sum of salicin, salicortin, homaloside D, and 6′-O-benzoylsalicortin), (b) salicortin; (c) 6′-O-benzoylsalicortin. (d) Flavan-3-ol catechin and (e) flavonoid rutin. All values are expressed as relative amounts, calculated as analyte peak area divided by internal standard peak area and normalized to sample dry weight (mg). For 6′-O-benzoylsalicortin and rutin, only leaf data are displayed as these compounds were not detected in the roots. Colours indicate water regime (field capacity: 100 %, blue; 50 %, beige; 25 % brown); striped bars denote herbivory. Linear mixed-effects models tested the effects of water regime, herbivory, and their interaction, including biomass dry weight as a covariate and genotype as a random effect (**Table S5**), (* p < 0.05; ** p < 0.01; ***p < 0.001; n.s. = not significant). Bars show means ± SE (n = 16-18). Note that the scaling of the y-axis differs between compounds.

### 3.3 Drought and herbivory increase phytohormone concentrations in *P. nigra* leaves

Phytohormone concentrations varied with water regime and herbivory treatments (**Fig. 3**; **Table S6**). Salicylic acid (SA) increased under drought conditions (**Fig. 3a**) and was significantly affected by water regime (p = 0.004), whereas herbivory had no significant effect. Abscisic acid (ABA) concentrations increased with decreasing water availability (**Fig. 3b**) and were significantly influenced by both water regime (p < 0.001) and herbivory (p = 0.012). Total jasmonates were consistently higher in herbivory-treated plants and increased with drought intensity but only within herbivore induced plants (**Fig. 3c**). Herbivory had a significant effect on total jasmonate levels (p < 0.001), whereas the main effect of water regime was not significant. However, a significant interaction between water regime and herbivory was detected (p < 0.001).

**Fig. 3:**
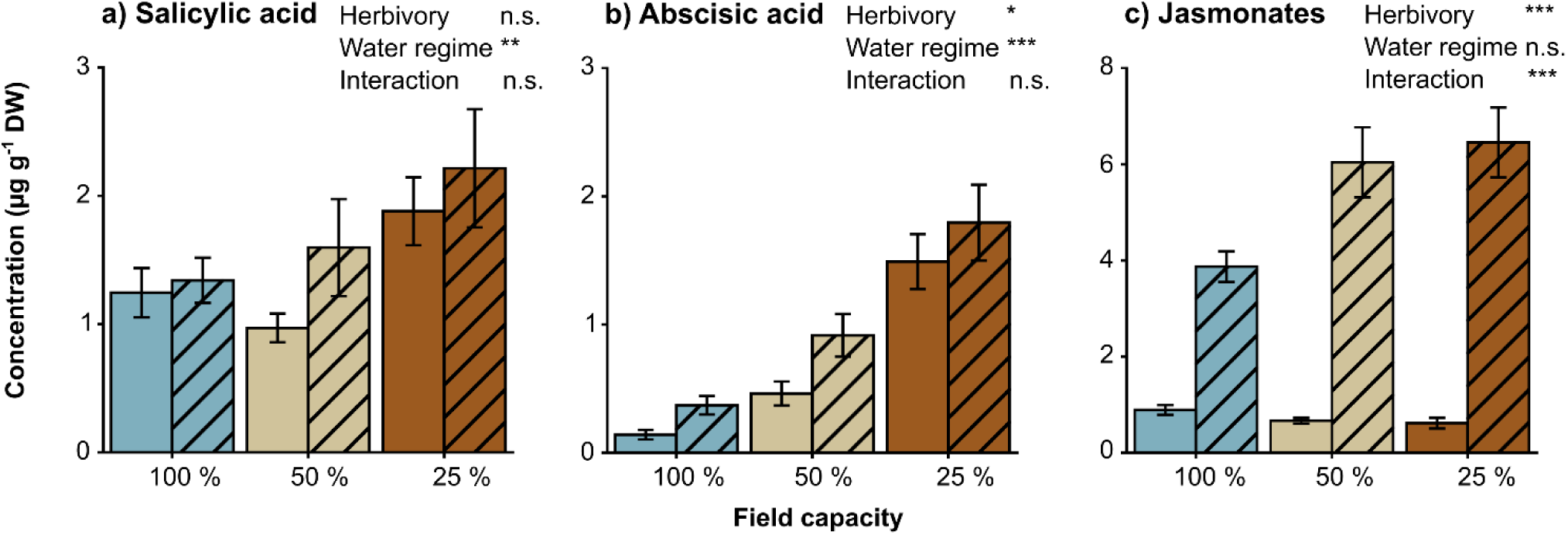
Effects of water regime and herbivory on phytohormones in *Populus nigra* leaves. Total jasmonates represent the sum of jasmonic acid (JA), (+)- and (-)-jasmonoyl isoleucine, hydroxy-JA, hydroxy-JA-Ile, carboxy-JA-Ile. Colours indicate water regime (field capacity: 100 %, blue; 50 %, beige; 25 % brown); striped bars denote the herbivory treatment with *Chrysomela tremulae* beetles. Linear mixed-effects models tested the effects of water regime, herbivory, and their interaction, including biomass dry weight as a covariate and genotype as a random effect (**Table S6**), (* p < 0.05; ** p < 0.01; *** p < 0.001; n.s. = not significant). Bars show means ± SE (n = 16-18).

### 3.4 Free amino acid profiles shift in response to water regime and herbivory

Relative changes in free amino acid concentrations compared with the constitutive well-watered control (field capacity 100 %) are shown in the heatmap **in Fig. 4c**. There are clear treatment-dependent shifts in amino acid profiles, primarily structured by water regime. Reduced water availability was associated with widespread changes in concentrations. In the absence of herbivory, amino acid concentrations declined consistently with increasing drought intensity. In contrast, herbivore-induced plants exhibited more heterogeneous pattern, including both increases and decreases. Response patterns differed among metabolites, reflecting varying sensitivity to drought and herbivory.

**Fig. 4:**
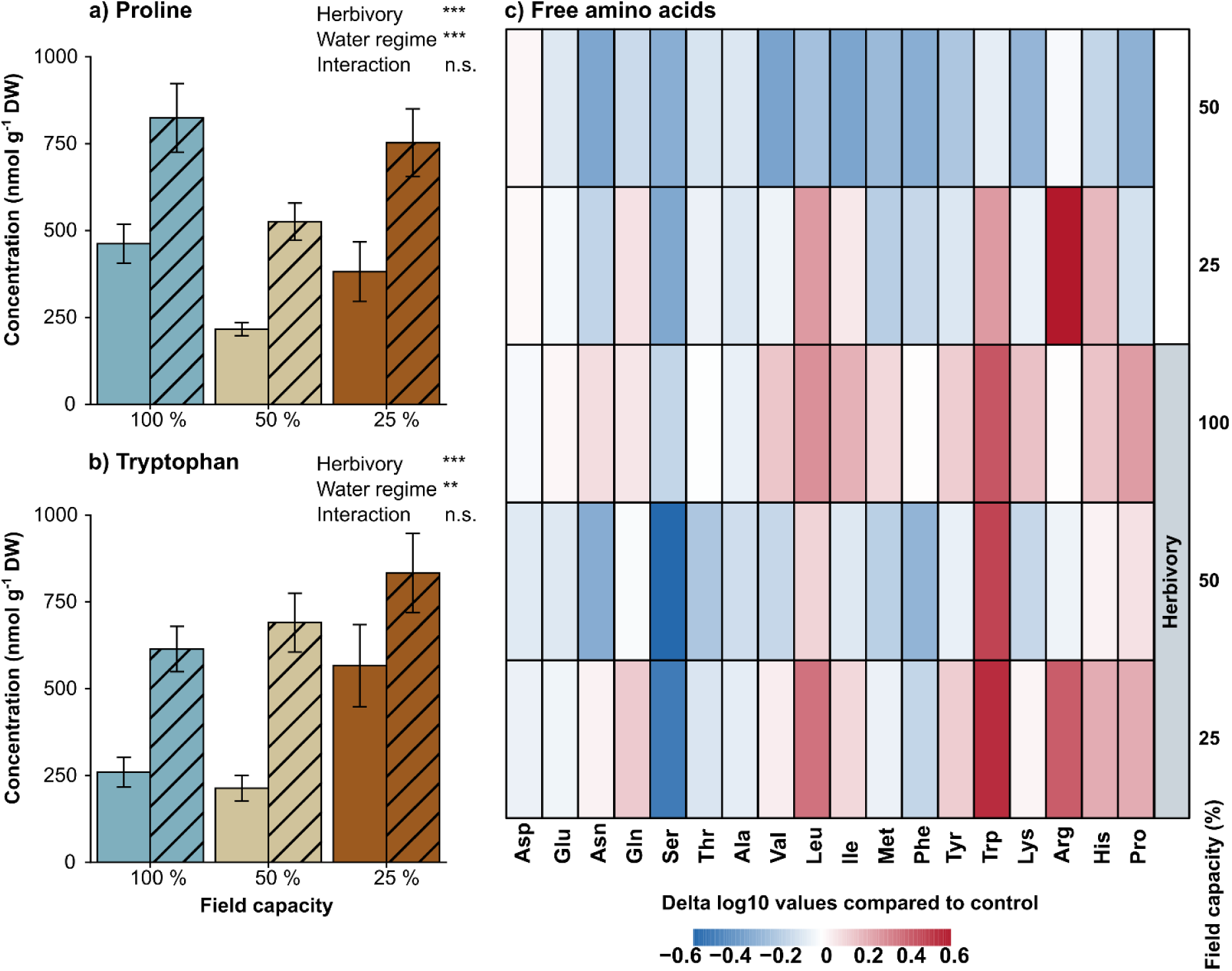
Effects of water regime and herbivory on free amino acids in *Populus nigra*: (a) Proline and (b) tryptophan concentrations, and (c) heatmap of delta log10 responses relative to the constitutive well-watered control. Amino acids in the heatmap are sorted according to their chemical classes. Colours indicate water regime (field capacity: 100 %, blue; 50 %, beige; 25 % brown); striped bars denote the herbivory treatment with *Chrysomela tremulae* beetles. Linear mixed-effects models tested the effects of water regime, herbivory, and their interaction, including biomass dry weight as a covariate and genotype as a random effect (**Table S8**), (* p < 0.05; ** p < 0.01; *** p < 0.001; n.s. = not significant). Bars show means ± SE (n = 16-18) (**Table S7**).

Concentrations of proline and tryptophan, selected based on the overall amino acid response patterns, are presented in **Fig. 4 a, b**. Proline concentrations were significantly influenced by both water regime and herbivory. Proline levels were highest in herbivore-induced plants under well-watered conditions (field capacity 100 %). Drought treatments altered proline concentrations, yet concentration remained consistently elevated in herbivore-induced compared with constitutive plants (p < 0.001). No significant water regime × herbivory interaction was detected. Tryptophan concentrations likewise varied with treatment. Both water regime and herbivory exerted significant main effects (p < 0.01), whereas their interaction was not statistically significant. Tryptophan concentrations were higher in herbivore-induced plants and varied across drought treatments with highest values in severe drought treatment. Full statistical analyses for all quantified free amino acids are provided in **Table S8.**

### 3.5 Water regime and herbivory strongly influence *P. nigra* VOC emission

Young *P. nigra* trees emitted VOC blends characterized by high chemical complexity, with 83 compounds detected in constitutive emissions and 86 compounds in trees suffering from beetle herbivory (**Table S10**). The detected VOCs belonged to several chemical classes, including monoterpenes, sesquiterpenes, homoterpenes, nitrogen-containing compounds, aromatics, green leaf volatiles (GLVs), and compounds that could not be assigned to any of these major groups and are thus termed “other compounds” from here on in the manuscript (**Table S9**).

Functional Hill diversity (q = 1) differed among treatments (**Fig. 5**). Statistical analysis revealed a significant interaction between water regime and herbivory (F = 6.45, p = 0.002; **Table S10**), while no significant main effects were observed. Across VOC classes, herbivory had a significant effect on emission rates (**Fig. 6; Table S11**), affecting both total VOC emissions and class-level emissions (p < 0.001). Drought significantly affected (*E*)-4,8-dimethyl-1,3,7-nonatriene (DMNT) emissions (F = 3.20, p = 0.045). With reduced water levels, DMNT emission decreased. No significant water regime effects or interaction was detected for other VOC classes. Although there were no significant effects of the water regime, trends can be observed in the response to water level, among the different volatile classes (**Fig. 6**). Under herbivory, total VOC emissions appeared highest under well-watered and moderate drought conditions, declining only under severe drought. In contrast, DMNT emissions showed a significant decrease with increasing drought intensity, and sesquiterpenes exhibited a similar pattern. Monoterpene emissions exhibited a stepwise pattern, remaining similarly high under well-watered and moderate drought conditions, followed by a reduction under severe drought. Conversely, nitrogen-containing compounds displayed an opposing trend, with emissions increasing under moderate and severe drought in herbivory-treated plants. GLVs displayed a similar drought-related pattern, with maximum emissions under severe drought (**Fig. 6**). However, these patterns were not statistically supported across VOC classes, with the exception of DMNT. Overall, herbivory emerged as the dominant trigger of VOC release, while water regime tends to influence the magnitude and compositional structure of induced blends in a class-specific way.

**Fig. 5:**
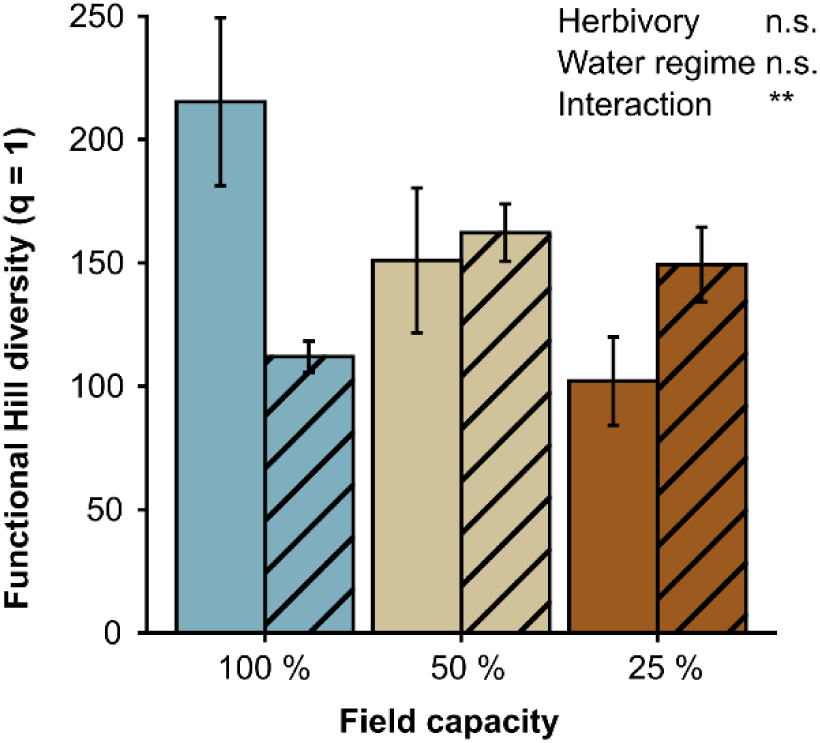
Functional Hill diversity (q₁) of VOC emissions under different drought and herbivory treatments in *Populus nigra*. q₁ reflects the effective diversity of common functional traits. Colours indicate water regime (field capacity: 100 %, blue; 50 %, beige; 25 % brown); striped bars denote the herbivory treatment with *Chrysomela tremulae* beetles. Linear mixed-effects models tested the effects of water regime, herbivory, and their interaction, including biomass dry weight as a covariate and genotype as a random effect (**Table S11**), (* p < 0.05; ** p < 0.01; *** p < 0.001; n.s. = not significant). Bars show means ± SE (n = 16-18).

**Fig. 6:**
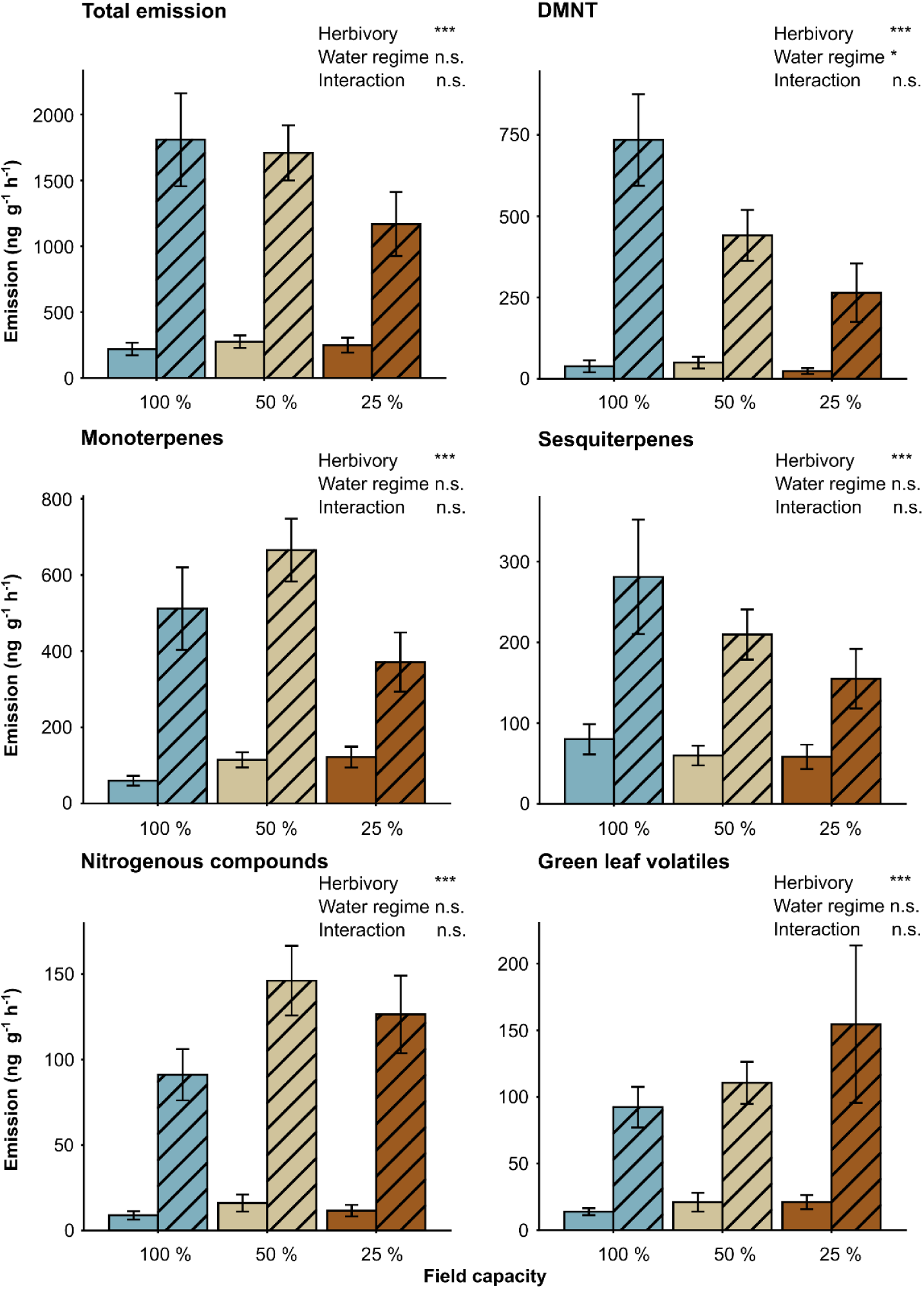
Effects of drought and herbivory on VOC emission by compound class in *Populus nigra*. Total VOC emission and emissions of the major VOC classes (monoterpenes, sesquiterpenes, nitrogenous compounds, green leaf volatiles, nitrogenous compounds) and DMNT across drought levels and herbivory treatments are shown. Colours indicate water regime (field capacity: 100 %, blue; 50 %, beige; 25 % brown); striped bars denote the herbivory treatment with *Chrysomela tremulae* beetles. Linear mixed-effects models tested the effects of water regime, herbivory, and their interaction, including biomass dry weight as a covariate and genotype as a random effect (**Table S11**), (* p < 0.05; ** p < 0.01; *** p < 0.001; n.s. = not significant). Bars show means ± SE (n = 16-18).

Treatment effects on the emission of individual VOCs are shown in **Table S12**. To illustrate compound-specific responses across different chemical classes, six representative VOCs were examined in more detail (**Fig. 7**), including the GLV 3-hexenal, the monoterpene (*E*)-β-ocimene, the aromatic compounds benzaldehyde and salicylaldehyde, the nitrogen-containing compound benzyl cyanide, and the monoterpene α-pinene. These compounds were chosen as representative VOCs because they are commonly reported herbivore-induced compounds and reflect different biosynthetic pathways. Across these compounds, herbivory had a significant main effect on all VOCs. Significant main effects of water regime were detected for all compounds except salicylaldehyde (p < 0.05). No statistically significant water regime × herbivory interactions were observed (p > 0.05) (**Fig.7**; **Table S12**).

**Fig. 7:**
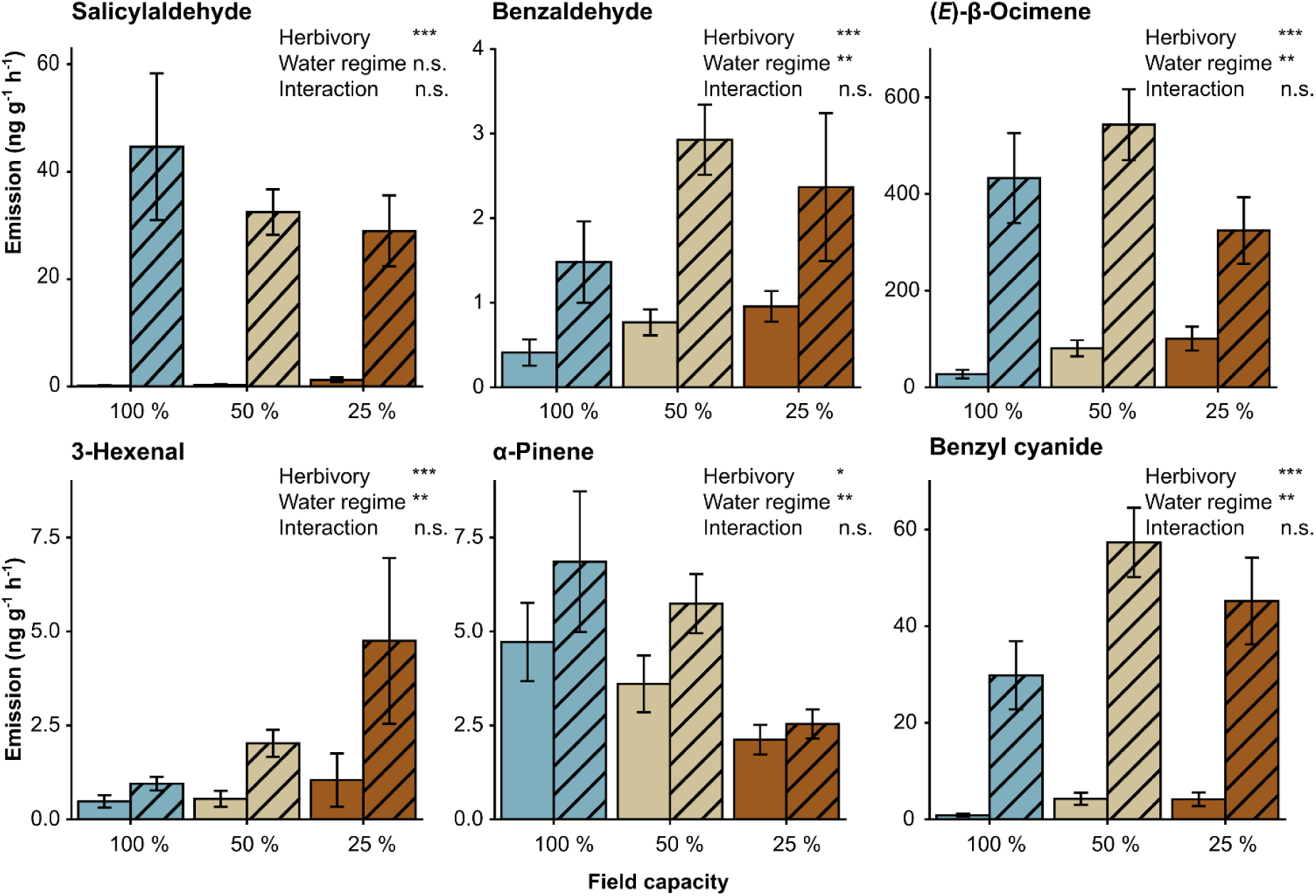
Compound-specific VOC emission responses to drought and herbivory in *Populus nigra*. Emission rates of selected volatile compounds (salicylaldehyde, benzaldehyde, (*E*)-β-ocimene, 3-hexenal, α-pinene, benzyl cyanide) are shown. Colours indicate water regime (field capacity: 100 %, blue; 50 %, beige; 25 % brown); striped bars denote the herbivory treatment with *Chrysomela tremulae* beetles. Linear mixed-effects models tested the effects of water regime, herbivory, and their interaction, including biomass dry weight as a covariate and genotype as a random effect (**Table S12**), (* p < 0.05; ** p < 0.01; *** p < 0.001; n.s. = not significant). Bars show means ± SE (n = 16-18).

These compounds exhibited distinct emission patterns across water regimes (**Fig. 7**; **Table S12**). Emissions of benzyl cyanide and (*E*)-β-ocimene, both characteristic herbivore-induced plant volatiles (HIPVs), increased strongly under herbivory, but showed distinct drought-dependent patterns. (*E*)-β-ocimene emission increased with drought intensity under constitutive conditions. Under herbivory, emission rates were highest under moderate drought, followed by a decline under severe drought, where values approached those observed under well-watered conditions. Benzyl cyanide emission likewise increased under drought in constitutive samples and herbivore-induced plants showed higher emission rates under drought treatments. 3-Hexenal emission increased with drought intensity, especially under herbivory, and was highest under severe drought conditions. Benzaldehyde emission increased under drought stress, both in constitutive and herbivore-induced samples, with a more pronounced response in herbivory-treated plants. In contrast, salicylaldehyde emission was primarily elevated under herbivory. α-Pinene emission declined with increasing drought intensity, indicating sensitivity to reduced water availability. Overall, emission responses differed markedly among compounds, demonstrating compound-specific effects of herbivory and drought.

To assess whether variation in VOC emissions was related to herbivore damage intensity, regression analyses were performed using herbivory-treated plants only. Within herbivore-induced plants, VOC emissions varied with herbivore damage intensity, indicating that emission rates were not solely determined by treatment effects but also by the extent of leaf damage. Across compounds, increasing damage was associated with compound-specific changes in VOC emission rates (**Table S13**). Several VOCs exhibited positive relationships with damage intensity, reflecting enhanced emissions at higher levels of herbivory, whereas others showed weaker or no significant associations (**Fig. S2**). Emission rates of multiple representative VOCs were significantly affected by leaf damage (p < 0.05). Significant positive relationships with damage intensity were detected for salicylaldehyde, (*E*)-β-ocimene, benzaldehyde, and benzyl cyanide, whereas α-pinene showed a weaker but still significant association. In contrast, 3-hexenal exhibited no significant damage effect. Under drought conditions, the relationship between herbivore damage and VOC emission rates was compound-specific. The strength of damage-emission relationships differed among water regime treatments for several compounds, with significant damage × water regime interactions detected for salicylaldehyde, benzaldehyde, (*E*)-β-ocimene and benzyl cyanide (p < 0.05). Across treatments, plants emitted a diverse spectrum of VOCs. Functional diversity, total emission rates, compound-class emissions, and individual VOC responses varied with water regime and herbivory. Herbivory consistently increased VOC emissions, whereas drought effects were compound- and class-specific. Compound-level linear mixed-effects models confirmed heterogeneous VOC responses, and regression analyses revealed compound-specific effects of herbivore damage intensity on emission rates.

## 4. Discussion

In this study, we tested how different water levels in combination with leaf chewing herbivory by the specialist beetle *C. tremulae* effect young *P. nigra* (black poplar) trees. Drought consistently shaped plant responses across morphological traits and several metabolic pathways, including phytohormones, amino acids, phenolics, and volatile organic compounds (VOCs). In leaves, water limitation significantly increased the concentration of the flavan-3-ol catechin and the flavonoid rutin, suggesting a general shift in secondary metabolism following abiotic stress (Holopainen et al., 2018; Kumar et al., 2023; Shomali et al., 2022). In roots of *P. nigra*, the concentrations of total salicinoids with salicortin as the major compound, decreased in response to severe water limitation, highlighting the importance of tissue-specific allocation of compounds when assessing plant stress responses (Holopainen et al., 2018). These drought-induced changes are likely to have important consequences for plant-herbivore interactions, given the well-established role of phenolic compounds, especially salicinoids, in the chemical defense of trees within the genus *Populus* (e.g. Boeckler et al., 2011; Osier and Lindroth, 2001; Kaufman et al., 2025). Notably, drought did not merely suppress or amplify herbivore-induced responses but rather altered the quantity and composition of induced chemical phenotypes. This pattern was most evident in the class- and compound-specific modulation of VOC profiles. Our results emphasize how abiotic stress can alter the chemical signalling and defense-related traits of trees, thereby modifying the ecological context of biotic interactions. In temperate forests, where droughts are becoming more frequent and severe due to climate change, these combinations of stressors will likely play an important role in shaping plant-environment interactions as well as the stability of forest food webs (Bansal et al., 2013; Holopainen et al., 2018; Malone et al., 2023).

Our data show that drought strongly affected morphological traits of young *P. nigra* trees like growth and biomass accumulation. Reduced growth is known to be one of the earliest and most consistent plant responses to water limitation, reflecting rapid physiological adjustments to declining water availability (Brunner et al., 2015; Mundim and Pringle, 2018). Such responses often involve reductions in height growth, biomass accumulation, and leaf area, which collectively limit transpiration and carbon demand under water deficit conditions (Günthardt-Goerg and Vollenweider, 2007; Rosso et al., 2023; Yin et al., 2004). Such early growth suppression is frequently interpreted as part of a drought-induced shift in resource allocation, whereby carbon investment into structural growth is reduced while resources are redirected towards stress tolerance and defense-related processes (Bogeat-Triboulot et al., 2007; Brunner et al., 2015; Mundim and Pringle, 2018). Experimental herbivory did not measurably affect growth-related traits in our study, which is certainly due to the relatively short feeding period. In woody plants, visible impacts of herbivory on growth often emerge only after prolonged or repeated feeding events that substantially alter carbon balance or tissue loss (Zvereva et al., 2012). Phytohormone and free amino acid levels in *P. nigra* responded strongly to increasing drought intensity. Notably, the substantial accumulation of ABA underscores its central role in early drought signaling, where it regulates stomatal closure and coordinates downstream transcriptional and metabolic responses, including modulation of root growth and resource allocation (Tiwari et al., 2017). Importantly, stomatal closure often happens early during drought adaptation to minimize water loss and keep up water potential (Schroeder et al., 2001; Waadt et al., 2022). With prolonged drought exposure, stomata closure lead to a decrease in carbon dioxide fixation, subsequently causing a reduction in photosynthesis and therefore growth (Niinemets, 2016; Ahmad et al., 2023; Ali et al., 2025; Rahmani et al., 2025; Zahedi et al., 2025). Additionally, an increase in jasmonates (jasmonic acid and its precursors) was observed in response to drought in the herbivore-treated *P. nigra* trees, a trend possibly due to the slightly higher herbivore damage in the water stressed trees. JA, but not SA levels increased in response to herbivory, consistent with previous findings in poplar by Boeckler et al. (2013) and Clavijo McCormick et al. (2019), and underscoring the central role of the JA pathway in herbivore-induced VOC emission (Kessler et al., 2004) and in regulating plant defense responses to insect herbivores, particularly chewing insects (Howe and Jander, 2008). JA and SA are often considered antagonistic, with SA suppressing JA accumulation (Berens et al., 2017). However, in *Populus*, their interaction appears less strictly antagonistic, as both JA- and SA- dependent pathways can be simultaneously upregulated in response to biotic stress such as fungal infection (Ullah et al., 2022). The concurrent increase in ABA and SA under drought suggests crosstalk between signaling pathways which have been described in both herbaceous and woody plant species (Fàbregas and Fernie, 2019; Huberty and Denno, 2004; Pieterse et al., 2012).

We observed pronounced shifts in amino acid profiles following herbivory, drought, and their combination, indicating rapid reconfiguration of primary metabolites in *P. nigra* in response to different stress conditions (Garavillon-Tournayre et al., 2018; Rosso et al., 2023). Amino acids can serve an important role in osmotic adjustment by helping to maintain homeostasis during drought stress, (Fàbregas and Fernie, 2019; Raza et al., 2026; Tschaplinski et al., 2019) but also as precursors for induced secondary metabolites (Heinemann and Hildebrandt, 2021; Zhou et al., 2015). For example, Alcázar et al. (2010) reported on the role of polyamines during abiotic stress, highlighting their accumulation as a conserved drought response and illustrating how amino acids such as arginine are not only involved in osmotic adjustment but also serve as precursors for metabolites that enhance stress tolerance. Although increased proline levels under drought have been reported (Bandurska et al., 2017; Barchet et al., 2014; Lei et al., 2007), accumulation typically occurs only under prolonged or severe water stress (Fàbregas and Fernie, 2019; Van Rensburg et al., 1993). Our results showed reduced proline levels under both 50 % and 25 % field capacity. Although proline is considered important for drought adaptation through osmotic adjustment and ROS scavenging, its regulation is complex (Verbruggen and Hermans, 2008). In *P. nigra*, proline increased in the herbivory treatment, where it could play a role in ROS buffering and stress signalling (Mithöfer and Boland, 2012; Szabados and Savouré, 2010). The decrease under drought hints at a possible metabolic impairment in its early stages. However, proline accumulation is driven by ABA among other factors (Hare et al., 1999; Yoshiba et al., 1997), which showed increased levels under drought conditions in our experiments. Depending on the stage of stress response the plants were in, the upregulation of proline might not have been induced yet, whereas earlier metabolic changes already happened (Fàbregas and Fernie, 2019; Hildebrandt, 2018). Tryptophan levels increased under drought, consistent with its role as a precursor of stress-related defense metabolites (Halkier and Gershenzon, 2006; Steinbrenner et al., 2011) and the general accumulation of aromatic amino acids during drought stress (Fàbregas and Fernie, 2019; Pires et al., 2016). In contrast to morphological changes or VOC emissions, the specific water stress level did markedly affect the amino acid response in *P. nigra*. While no increase could be observed in plants exposed to moderate levels of water deficit, the concentrations of several amino acids were elevated in plants kept at only 25 % field capacity without herbivory exposure. This is in line with results shown by Todaka et al., (2017), who could observe significant increase of amino acid levels in rice plants in response to severe drought stress. However, drought treatment leading to these changes was likely much more severe there, as leaves already wilted under severe drought conditions in this study. A general rise of amino acids following dry conditions has also been reported in poplar by Lackner et al. (2019), who focussed on belowground herbivory inevitably affecting root growth and therefore water availability. Since we observed changes in only a few amino acids (Trp, Arg, His, Leu), it seems likely that these are among the most drought-sensitive amino acids that increase early during drought response. These metabolic processes involving amino acids may represent the first stage of drought acclimation, enabling plants to maintain cellular homeostasis in the face of declining water availability (Hildebrandt, 2018; Shinozaki and Yamaguchi-Shinozaki, 2007). In herbivore-exposed plants, levels of several amino acids rose, which further supports the suggestion of shift from growth to defense in reaction to stressors (Herms and Mattson, 1992).

In contrast to leaf phytohormones and amino acids, our data show that phenolic compounds in roots exhibited more selective responses. Salicinoids, the major group of phenolic anti-herbivore defenses in poplar, showed only minor changes in response to drought, herbivory, or their combination. Similar patterns of relatively stable salicinoid levels under short-term environmental stress have been reported in previous studies (Boeckler et al. 2013; Holopainen et al., 2018). A notable exception to this relatively conservative response marked catechin, the monomer of condensed tannins (Barbehenn and Constabel, 2011; Boeckler et al., 2013), the second major group of poplar phenolics. Under drought stress, the relative amounts of catechin in *P. nigra* leaves increased significantly, which has been reported before for *Populus* (Barchet et al., 2014); however, in roots, it declined with increasing drought stress. The contrasting patterns observed in above- and belowground tissues suggest that drought can redirect phenolic metabolism in different ways (Lackner et al., 2019), potentially reflecting changes in carbon allocation and not only compound-, but organ-specific defense priorities (Yepes-Vivas et al., 2025). Phenolic compounds may accumulate during drought to support ROS scavenging and tissue protection (Hura et al., 2008) although the flavonoid rutin increased in leaves but declined in roots.

Herbivory acted as the primary trigger of VOC emissions throughout all investigated compound classes. These results are consistent with the role of VOCs as rapid, on-demand signals in plant-insect interactions (Clavijo McCormick et al., 2014b; Eberl et al., 2018; Unsicker et al., 2009). Notably, emission increased with feeding area across different compounds as it was also shown for a few *P. nigra* VOCs emitted by old-growth trees (Clavijo McCormick et al., 2019). This is an indicator for functionality of HIPVs that serve different roles in plant defense other than deterrence of herbivores. (*E*)-β-ocimene and DMNT both showed a positive correlation between feeding area and volatile emission and are known for playing a role in tri-trophic defense by attracting enemies of plant herbivores (Arimura et al., 2009; Dicke et al., 2009; Unsicker et al., 2009). The same pattern could be observed for benzaldehyde and benzyl cyanide. Although they were emitted in much smaller quantities, they still played an important role in differentiating blends from well-watered and drought-exposed plants. Since they reflect both defense signal and physiological plant status, they may function as “honest signals”. Drought alone had little effect on the levels of constitutive VOC emission, and most compound-classes were not drought-sensitive, except for the homoterpene DMNT. However, changes in composition and quantity of herbivore-induced VOCs could be observed, but in a highly compound- and class-specific manner instead of a general pattern.

Herbivory x water regime interaction did not lead to significant changes in VOC emissions. However, a trend for decreasing emissions under combined herbivory and drought stress could be observed for mono- and sesquiterpene emissions and, since terpenes make up a significant portion of volatile emissions in poplar, total emissions. Reduction in isoprene emission in response to a decrease in carbon availability because of drought exposure has been proposed before (Lavoir et al., 2009; Niinemets et al., 2010), although responses are usually complex. Studies on *Arabidopsis* could show that for short-term water limitations, there is no carbon starvation as plant growth is inhibited before photosynthesis, but instead the carbon could be allocated towards defense compound synthesis. For DMNT, Holopainen and Gershenzon (2010) reported increasing emissions in response to drought, whereas in our results, there was a significant decrease in response to drought and a decreasing trend for interaction. The combination of abiotic and biotic stress could exceed the response threshold, slowly leading to an increase of oxidative stress and less carbon gain. This is supported by the functional Hill diversity that showed a significant effect of drought x herbivory interaction on the compound functional diversity, hinting at a cross-talk between herbivory and drought defense pathways (Xiao et al., 2025). The functional Hill diversity was chosen as the preferred approach because it highlights variation in structural composition of VOC profiles. Structurally distinct compounds differ substantially in their ecological functions, e. g. by mediating herbivore interactions (Petrén et al., 2024). Functional diversity was higher in plants exposed to herbivory and water stress when compared with well-watered, herbivory-exposed plants. This suggests that drought may act as a modifier of HIPVs (Malone et al., 2023; Scott et al., 2019), although in order to test that hypothesis further, the severity and duration of drought stress needs to be adjusted.

Drought did not uniformly suppress metabolic activity in young *P. nigra* trees. While growth was strongly inhibited, constitutive VOC emissions remained largely stable under drought alone and declined primarily under the combined pressure of drought and herbivory. This pattern is consistent with the framework proposed by Hummel et al. (2010), suggesting that carbon starvation is not an inevitable consequence of drought since growth is inhibited more strongly than photosynthesis, leading to a more favorable C balance since growth is reduced but metabolic pathways are maintained. Plants may therefore maintain metabolic pathways while reallocating carbon resources. Earlier studies have reported contrasting responses. Niinemets (2010) emphasized that even mild drought may reduce photosynthetic carbon assimilation and affect volatile emissions. Similarly, Gouinguené and Turlings (2002) observed increased HIPV emissions under drought in maize, including enhanced DMNT release. In contrast, we detected reduced DMNT emissions. These discrepancies likely reflect species-specific physiological strategies and environmental context. Compound-class-specific responses further underline the complexity of drought effects. Terpene emissions tended to decrease under combined stress, whereas GLV emissions showed a tendency to increase, consistent with rapid activation of the LOX pathway under abiotic stress (Wang et al., 2021). Random Forest analyses and modeling approaches additionally highlighted nitrogen-containing compounds such as aldoximes, which are known to contribute to plant defense (Clavijo Mccormick et al., 2014b; Douma et al., 2019; Irmisch et al., 2013). However, interpretation of drought-related effects should consider several constraints. While it is possible to get an overview of the shifts in plant metabolism and emissions based on the results that were obtained, supplementary measurements of the plants physiological state, stomatal conductance, leaf water potential and photosynthetic capacity in particular, could help understand adaptation mechanisms. Although our result did not show a drastic change in response between 50 % and 25 % field capacity, plant response is likely still influenced by the severity of drought stress (Fàbregas and Fernie, 2019; Niinemets, 2010). The drought application time of five days at required field capacity is on the shorter end of drought exposure, which may have had consequences for the perceived severity of drought stress and photosynthetic impairment.

Our study focuses on young trees, which may differ considerably from mature individuals in their responses to drought and herbivory. Juvenile trees typically show higher growth rates and metabolic activity but are also more sensitive to water limitation (Niinemets, 2002; Thomas, 2010). Mature trees, by contrast, often benefit from larger carbon reserves and more developed hydraulic systems, which can buffer stress effects (Au et al., 2022; Bennett et al., 2015). Ontogenetic changes in physiology and defense have been described across a range of species, including *Acer*, *Fraxinus*, and *Quercus* (Niinemets, 2002; Thomas, 2010). This is particularly relevant in the context of global change, as large trees dominate carbon storage and ecosystem functioning. The extent to which the patterns observed here translate to mature trees therefore remains an open question and should be addressed in future studies.

In summary, our results highlight the complex nature of plant responses to interacting abiotic and biotic stressors. Drought acted as a primary driver of plant phenotype by strongly inhibiting growth, while its effects on chemical traits and VOC emissions were predominantly compound-and compound class-specific rather than uniformly suppressive. Herbivory emerged as the dominant trigger of VOC emissions, with drought modulating the magnitude and composition of selected responses. Although water regime x herbivory interactions were less pronounced than expected, the predominantly additive patterns observed across traits underscore the stability and flexibility of defense-related metabolic processes. Together, these findings demonstrate that moderate drought primarily constrains growth while allowing defense-associated pathways to remain functional and dynamically regulated.

## 6. Acknowledgements

We want to thank Jonathan Gershenzon for providing the analytics for phytohormone and amino acid analysis. We thank Liv Kraack, Gunda M. Teichmann, Elina Negwer, Sirui Xing and Frauke Harmel for their support in various aspects of tree cultivation, beetle rearing and plant harvesting. We thank Beate Rothe for preparing the samples for phytohormone and amino acid analysis. We are grateful to Christin Walther and Pamela Medina van Berkum for advice on analytical procedures and statistics. A special thanks to Alexandra Erfmeier for providing us with greenhouse space for our experiment.

## 8. Contributions

SKW, HG and SBU conceived the project. J-PS provided the *P. nigra* genotypes. SKW and HG did the sampling and analysed the data. SKW, HG and MR performed the chemical analyses. SKW and HG wrote the paper. J-PS, EK and SBU edited the manuscript and all authors reviewed the manuscript.

## 9. Supporting Information

**Fig. S1**. Overview of the experimental design.

**Fig. S2.** Relationship between herbivore damage intensity and VOC emission rates in herbivory-treated plants.

**Table S1.** Overview of experimental treatments and water regime conditions.

**Table S2.** Parameters used in LC-MS/MS analysis of phytohormones.

**Table S3.** Parameters used in LC-MS/MS analysis of amino acids.

**Table S4.** Statistical results for linear mixed-effects models testing the effects of water regime, herbivory, and their interaction on morphological traits.

**Table S5.** Statistical results for linear mixed-effects models testing the effects of water regime, herbivory, and their interaction on phenolic compound concentrations in leaves and roots.

**Table S6.** Statistical results for linear mixed-effects models testing the effects of water regime, herbivory, and their interaction on phytohormone concentrations.

**Table S7.** Mean concentrations of individual free amino acids under different water regime and herbivory treatments.

**Table S8.** Statistical results for linear mixed-effects models testing the effects of water regime, herbivory, and their interaction on free amino acid concentrations.

**Table S9.** Overview of all identified volatile compounds.

**Table S10.** Sample sizes and number of detected volatile organic compounds (VOCs) across treatments.

**Table S11.** Statistical results for linear mixed-effects models testing the effects of water regime, herbivory, and their interaction on total VOC emissions, VOC classes, and functional hill diversity (q₁).

**Table S12.** Statistical results for linear mixed-effects models testing the effects of water regime, herbivory, and their interaction on selected volatile organic compounds.

**Table S13.** Statistical results for linear mixed-effects regression models testing the effects of leaf damage intensity, water regime, and their interaction on individual VOC emissions.

