## Supplemental material for "Drought and Herbivory Shape Morphological and Chemical Traits in Black Poplar (*Populus nigra*)"

**Fig. S1**. Overview of the experimental design.

**Fig. S2.** Relationship between herbivore damage intensity and VOC emission rates in herbivory-treated plants.

**Table S1.** Overview of experimental treatments and water regime conditions.

**Table S2.** Parameters used in LC-MS/MS analysis of phytohormones.

**Table S3.** Parameters used in LC-MS/MS analysis of amino acids.

**Table S4.** Statistical results for linear mixed-effects models testing the effects of water regime, herbivory, and their interaction on morphological traits.

**Table S5.** Statistical results for linear mixed-effects models testing the effects of water regime, herbivory, and their interaction on phenolic compound concentrations in leaves and roots.

**Table S6.** Statistical results for linear mixed-effects models testing the effects of water regime, herbivory, and their interaction on phytohormone concentrations.

**Table S7.** Mean concentrations of individual free amino acids under different water regime and herbivory treatments.

**Table S8.** Statistical results for linear mixed-effects models testing the effects of water regime, herbivory, and their interaction on free amino acid concentrations.

**Table S9.** Overview of all identified volatile compounds.

**Table S10.** Sample sizes and number of detected volatile organic compounds (VOCs) across treatments.

**Table S11.** Statistical results for linear mixed-effects models testing the effects of water regime, herbivory, and their interaction on total VOC emissions, VOC classes, and functional hill diversity ($q_1$).

**Table S12.** Statistical results for linear mixed-effects models testing the effects of water regime, herbivory, and their interaction on selected volatile organic compounds.

**Table S13.** Statistical results for linear mixed-effects regression models testing the effects of leaf damage intensity, water regime, and their interaction on individual VOC emissions.

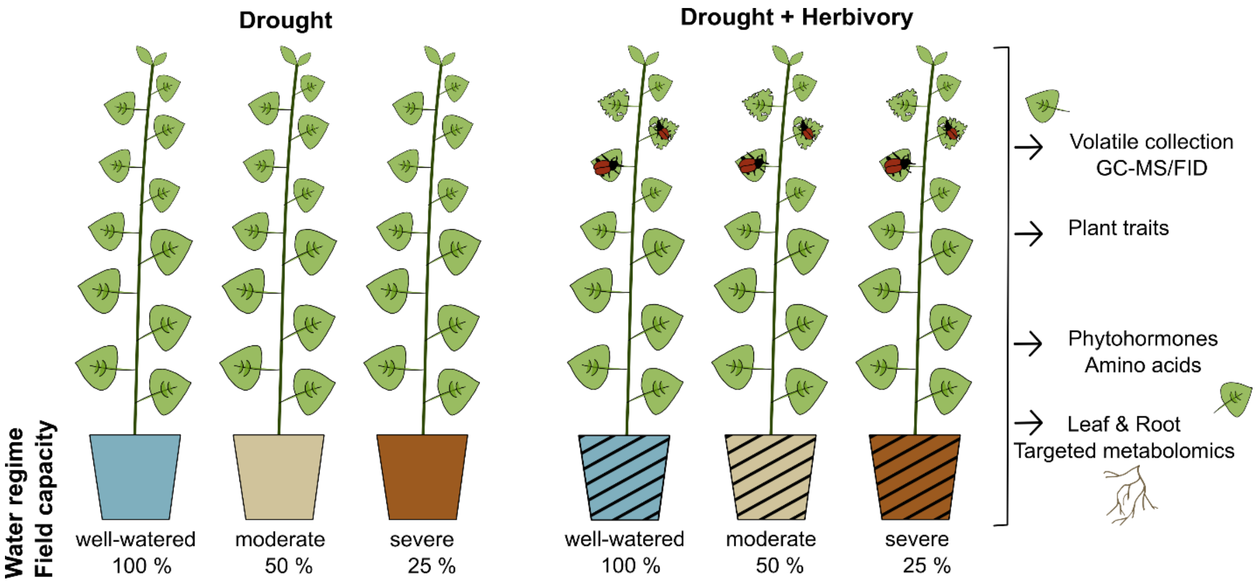


**Fig. S1. Graphical illustration of the experimental design.** Young *Populus nigra* trees were exposed to three water regime treatments corresponding to defined field capacity levels: well-watered (FC 100%), moderate drought (FC 50%), and severe drought (FC 25%). Drought conditions were maintained for five days following stress establishment. During the final two days, plants from three of the six experimental groups were subjected to herbivory. Colours indicate water regime (field capacity); striped pots denote herbivory. The experiment included three genotypes, with 4-6 plants per treatment combination and genotype. Morphological traits, volatile organic compounds (VOCs), phenolic compounds, phytohormones, and free amino acids were quantified.

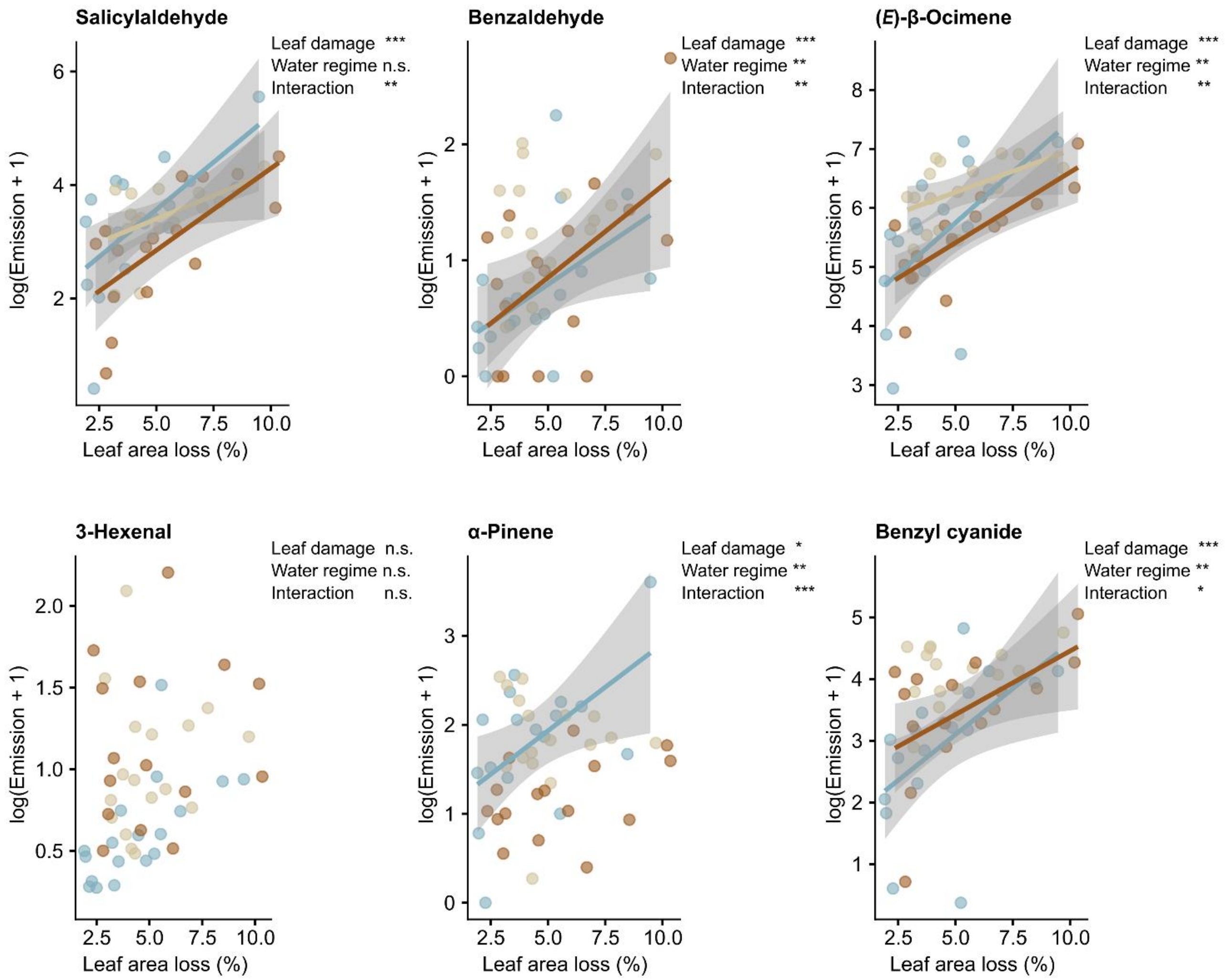


**Fig. S2. Relationship between herbivore damage intensity and VOC emission rates in herbivory-treated plants.** Points represent individual biological replicates. Colours indicate water regime treatments (blue: well-watered (FC 100%), beige: moderate drought (FC 50%), brown: severe drought (FC 25%)). Lines show linear regressions fitted separately for each water regime and are displayed only when the damage effect was statistically significant. Shaded areas represent 95% confidence intervals. VOC emission rates were natural log-transformed (ln(x + 1)) prior to analysis. Statistical analyses were performed using linear models (LMs), including leaf area loss (%), water regime, and their interaction as fixed effects (Table S13). Significance levels: * $p < 0.05$; ** $p < 0.01$; *** $p < 0.001$; n.s. = not significant.

**Table S1. Overview of experimental treatments and water regime conditions.**
The table summarizes treatment combinations, sample sizes (n), and corresponding water availability levels. Water regimes are expressed as percentages of field capacity and associated soil moisture values. Abbreviations: Field capacity (FC, %); Soil moisture (SM, volumetric %).

| Water regime | Herbivory | Replicates | FC (%) / SM (%) |
|---|---|---|---|
| well-watered (0) | no | 18 (m25=7, m30=5, m32=6) | 100 / 50 - 60 |
| moderate (1) | no | 18 (m25=6, m30=6, m32=6) | 50 / 15 - 20 |
| severe (2) | no | 18 (m25=6, m30=6, m32=6) | < 25 / < 15 |
| well-watered (0) | yes | 18 (m25=7, m30=5, m32=6) | 100 / 50 - 60 |
| moderate (1) | yes | 18 (m25=6, m30=5, m32=7) | 50 / 15 - 20 |
| severe (2) | yes | 18 (m25=6, m30=6, m32=4) | < 25 / < 15 |

**Table S2. Parameters for LC-MS/MS analysis of phytohormones.** Analyses were performed on a triple quadrupole instrument (HPLC 1260 (Agilent Technologies)-QTRAP6500 (SCIEX)).

| Compound | Q1 (Da) | Q3 (Da) | RT (min) | Internal standard | DP (V) | CE (V) |
|---|---|---|---|---|---|---|
| SA | 136.93 | 93 | 3.3 | D4-SA | -20 | -24 |
| ABA | 263 | 153.2 | 3.4 | D6-ABA | -20 | -22 |
| JA | 209.07 | 59 | 3.6 | D6-JA | -20 | -24 |
| JA-Ile | 322.19 | 130.1 | 3.9 | D6-JA-Ile | -20 | -30 |
| OH-JA-Ile | 338.1 | 130.1 | 3.0 | D6-JA-Ile | -20 | -30 |
| OH-JA | 225.1 | 59 | 2.6 | D6-JA | -20 | -24 |
| COOH-JA-Ile | 352.1 | 130.1 | 3.0 | D6-JA-Ile | -20 | -30 |
| SA-Glucoside | 299.13 | 136.9 | 1.8 | D4-SA | -20 | -18 |
| D4-SA | 140.93 | 97 | 3.3 | | -20 | -24 |
| D6-ABA | 269 | 159.2 | 3.4 | | -20 | -22 |
| D6-JA | 215 | 59 | 3.6 | | -20 | -24 |
| D5-JA | 214 | 59 | 3.6 | | -20 | -24 |
| D6-JA-Ile | 328.19 | 130.1 | 3.9 | | -20 | -30 |
| D5-JA-Ile | 327.19 | 130.1 | 3.9 | | -20 | -30 |

**Abbreviations:** Q1, quadrupole 1; Q3, quadrupole 3; RT, retention time; DP, declustering potential; CE, collision energy. SA, salicylic acid; ABA, abscisic acid; JA, jasmonic acid; JA-Ile, jasmonoyl-isoleucine; OH-JA-Ile, hydroxy-jasmonoyl-isoleucine; OH-JA, hydroxy-jasmonic acid; COOH-JA-Ile, carboxy-jasmonoyl-isoleucine; SA-Glucoside, salicylic acid glucoside.

**Table S3. Parameters for LC-MS/MS analysis of amino acids.** Analyses were performed on a triple quadrupole instrument (HPLC 1260 (Agilent Technologies)-QTRAP6500 (SCIEX)).

| Compound | Q1 (Da) | Q3 (Da) | RT (min) | Internal standard | IS Q1 | IS Q3 | DP (V) | CE (V) |
|---|---|---|---|---|---|---|---|---|
| Ala | 90.1 | 44.1 | 0.5 | 13C,15N-Ala | 94.1 | 47.1 | 20 | 17 |
| Ser | 106.0 | 60.1 | 0.5 | 13C,15N-Ser | 110.0 | 63.1 | 20 | 15 |
| Pro | 116.1 | 70 | 0.5 | 13C,15N-Pro | 122.1 | 75.0 | 20 | 19 |
| Val | 118.1 | 72.2 | 0.5 | 13C,15N-Val | 124.1 | 77.2 | 20 | 13 |
| Thr | 120.1 | 74.2 | 0.5 | 13C,15N-Thr | 125.1 | 78.2 | 20 | 13 |
| Ile | 132.2 | 86.1 | 1.1 | 13C,15N-Ile | 139.2 | 92.1 | 20 | 13 |
| Leu | 132.2 | 86.1 | 1.3 | 13C,15N-Leu | 139.2 | 92.1 | 20 | 13 |
| Asp | 134.1 | 74.1 | 0.5 | 13C,15N-Asp | 139.1 | 77.1 | 20 | 19 |
| Glu | 148.1 | 102.1 | 0.5 | 13C,15N-Glu | 154.1 | 107.1 | 20 | 15 |
| Met | 150.2 | 104.1 | 0.7 | 13C,15N-Met | 156.2 | 109.1 | 20 | 13 |
| His | 156.2 | 110.1 | 0.4 | 13C,15N-His | 165.2 | 118.1 | 20 | 17 |
| Phe | 166.2 | 120.2 | 2.6 | 13C,15N-Phe | 176.2 | 129.2 | 20 | 17 |
| Arg | 175.1 | 70.1 | 0.4 | 13C,15N-Arg | 185.1 | 75.1 | 20 | 31 |
| Tyr | 182.1 | 136.2 | 1.4 | 13C,15N-Tyr | 192.1 | 145.2 | 20 | 17 |
| Asn | 133.1 | 74.1 | 0.5 | 13C,15N-Asp | | | 20 | 21 |
| Gln | 147.1 | 130 | 0.5 | 13C,15N-Gln | 154.1 | 136.0 | 20 | 13 |
| Trp | 205.2 | 188.1 | 3.2 | D5-Trp | 210.0 | 193.0 | 20 | 13 |

**Abbreviations:** Q1, quadrupole 1; Q3, quadrupole 3; RT, retention time; DP, declustering potential; CE, collision energy. Ala, alanine; Asp, aspartic acid; Asn, asparagine; Glu, glutamic acid; Gln, glutamine; His, histidine; Ile, isoleucine; Leu, leucine; Met, methionine; Phe, phenylalanine; Pro, proline; Ser, serine; Thr, threonine; Trp, tryptophan; Tyr, tyrosine; Val, valine.

**Table S4. Statistical results for linear mixed-effects models testing the effects of water regime, herbivory, and their interaction on morphological traits of *Populus nigra* leaves.** Response variables included height increase, biomass dry weight, and herbivory-induced leaf area loss (% of total leaf area). Linear mixed-effects models included water regime, herbivory, and their interaction as fixed effects, and genotype as a random effect. Type III ANOVA was applied. Degrees of freedom were estimated using the Satterthwaite approximation. The table reports F-values and p-values, marginal and conditional coefficients of determination ($R^2m/R^2c$), and the transformations applied to each response variable. Significant effects are shown in bold ($p < 0.05$). Sample size: n = 16-18.

| | | **Water regime** | | **Herbivory** | | **Interaction** | | |
|---|---|---|---|---|---|---|---|---|
| **Response** | **Transformation** | **F-statistic** | **p-value** | **F-statistic** | **p-value** | **F-statistic** | **p-value** | **$R^2m/R^2c$** |
| Height increase | log | 33.14 | **0.000** | 0.06 | 0.814 | 0.69 | 0.502 | 0.383 / 0.411 |
| Biomass (DW) | none | 3.49 | **0.034** | 0.03 | 0.866 | 0.08 | 0.927 | 0.057 / 0.168 |
| Leaf area loss | none | 1.81 | 0.174 | | | | | 0.049 / 0.292 |

**Table S5. Statistical results for linear mixed-effects models testing the effects of water regime, herbivory, and their interaction on phenolic compound concentrations in leaves and roots of *Populus nigra*.** Mixed-effects models included water regime, herbivory, and their interaction as fixed effects, leaf biomass dry weight as a covariate, and genotype as a random effect. Type III ANOVA was applied. Degrees of freedom were estimated using the Satterthwaite approximation. The table reports F-values and p-values, marginal and conditional coefficients of determination ($R^2m/R^2c$), and the transformations applied to each response variable. Significant effects are shown in bold ($p < 0.05$). Sample size: n = 16-18.

| | | Water regime | | Herbivory | | Interaction | | |
|---|---|---|---|---|---|---|---|---|
| **Response** | **Transformation** | **F-statistic** | **p-value** | **F-statistic** | **p-value** | **F-statistic** | **p-value** | **$R^2m/R^2c$** |
| **Leaves** | | | | | | | | |
| Total salicinoids | none | 2.26 | 0.109 | 6.30 | **0.014** | 2.88 | 0.061 | 0.10 / 0.39 |
| Salicortin | none | 2.68 | 0.074 | 3.80 | 0.054 | 2.09 | 0.128 | 0.09 / 0.34 |
| Homaloside D | none | 4.03 | **0.021** | 5.78 | **0.018** | 1.47 | 0.235 | 0.11 / 0.38 |
| Salicin | sqrt | 2.17 | 0.119 | 1.71 | 0.194 | 0.57 | 0.568 | 0.05 / 0.23 |
| 6`-O-benzoylsalicortin | none | 7.05 | **0.001** | 4.95 | **0.028** | 4.36 | **0.015** | 0.10 / 0.68 |
| Catechin | log | 17.66 | **0.000** | 3.76 | 0.055 | 0.22 | 0.803 | 0.25 / 0.34 |
| Rutin | sqrt | 3.21 | **0.044** | 5.38 | **0.022** | 0.11 | 0.898 | 0.06 / 0.52 |
| **Roots** | | | | | | | | |
| Total salicinoids | sqrt | 4.27 | **0.017** | 0.03 | 0.868 | 0.01 | 0.990 | 0.16 / 0.25 |
| Salicortin | sqrt | 4.96 | **0.009** | 0.08 | 0.781 | 0.04 | 0.965 | 0.14 / 0.25 |
| Homaloside D | sqrt | 4.07 | **0.020** | 0.03 | 0.871 | 0.09 | 0.913 | 0.17 / 0.24 |
| Catechin | sqrt | 5.15 | **0.007** | 0.25 | 0.617 | 1.42 | 0.246 | 0.12 / NA |

**Table S6. Statistical results for linear mixed-effects models testing the effects of water regime, herbivory, and their interaction on phytohormone concentrations in leaves of *Populus nigra*.** Mixed-effects models included water regime, herbivory, and their interaction as fixed effects, and genotype as a random effect. Type III ANOVA was applied. Degrees of freedom were estimated using the Satterthwaite approximation. The table reports F-values and p-values, marginal and conditional coefficients of determination ($R^2m/R^2c$), and the transformations applied to each response variable. Significant effects are shown in bold ($p < 0.05$). Sample size: n = 16-18.

| | | **Water regime** | | **Herbivory** | | **Interaction** | | |
|---|---|---|---|---|---|---|---|---|
| **Response** | **Transformation** | **F-statistic** | **p-value** | **F-statistic** | **p-value** | **F-statistic** | **p-value** | **$R^2m/R^2c$** |
| ABA | none | 39.01 | **0** | 6.56 | **0.012** | 0.28 | 0.753 | 0.43 / 0.46 |
| COOH-JA-Ile | none | 1.56 | 0.214 | 58.62 | **0** | 0.64 | 0.532 | 0.33 / 0.46 |
| JA | none | 0.38 | 0.683 | 55.33 | **0** | 3.55 | **0.032** | 0.36 / 0.40 |
| JA-Ile | none | 3.69 | **0.028** | 92.18 | **0** | 9.65 | **0** | 0.52 / 0.53 |
| total Jasmonates | sqrt | 2.12 | 0.14 | 360.67 | **0** | 8.24 | **0** | 0.78 / 0.78 |
| OH-JA | sqrt | 4.98 | **0.009** | 362.5 | **0** | 6.64 | **0.002** | 0.75 / 0.79 |
| OH-JA-Ile | none | 3.3 | **0.041** | 182.67 | **0** | 2.72 | 0.071 | 0.62 / 0.67 |
| OPDA | sqrt | 12.51 | **0** | 34.46 | **0** | 0.61 | 0.543 | 0.37 / 0.39 |
| SA | sqrt | 5.94 | **0.004** | 2 | 0.16 | 0.54 | 0.587 | 0.11 / 0.20 |
| SA-Glucoside | log | 1.09 | 0.339 | 0.11 | 0.745 | 0.44 | 0.647 | 0.02 / 0.20 |

**Abbreviations:** ABA, abscisic acid; COOH-JA-Ile, carboxy-jasmonoyl-isoleucine; JA, jasmonic acid; JA-Ile, jasmonoyl-isoleucine; total jasmonates, sum of quantified jasmonate-related compounds; OH-JA, hydroxy-jasmonic acid; OH-JA-Ile, hydroxy-jasmonoyl-isoleucine; OPDA, 12-oxo-phytodienoic acid; SA, salicylic acid; SA-Glucoside, salicylic acid glucoside

**Table S7. Mean concentrations of individual free amino acids under different water regime and herbivory treatments.** Standard errors and sample sizes (n) are provided.

| Amino acid | Water regime | Herbivory | n | Mean (nmol $g^{-1}$ DW) | SE (nmol $g^{-1}$ DW) |
|---|---|---|---|---|---|
| Ala | well-watered | minus | 18 | 3185.14 | 279.01 |
| Ala | moderate | minus | 17 | 2567.13 | 221.94 |
| Ala | severe | minus | 18 | 2533.85 | 177.05 |
| Ala | well-watered | plus | 18 | 2674.26 | 117.37 |
| Ala | moderate | plus | 19 | 2269.96 | 107.55 |
| Ala | severe | plus | 16 | 2645.12 | 192.13 |
| Arg | well-watered | minus | 18 | 169.03 | 63.84 |
| Arg | moderate | minus | 17 | 126.40 | 17.73 |
| Arg | severe | minus | 18 | 749.65 | 151.75 |
| Arg | well-watered | plus | 18 | 148.18 | 37.79 |
| Arg | moderate | plus | 19 | 148.37 | 35.09 |
| Arg | severe | plus | 16 | 418.19 | 103.67 |
| Asn | well-watered | minus | 18 | 880.46 | 134.28 |
| Asn | moderate | minus | 17 | 418.00 | 75.94 |
| Asn | severe | minus | 18 | 619.60 | 109.39 |
| Asn | well-watered | plus | 18 | 1215.71 | 247.04 |
| Asn | moderate | plus | 19 | 446.32 | 86.29 |
| Asn | severe | plus | 16 | 1026.67 | 218.14 |
| Asp | well-watered | minus | 18 | 3886.76 | 344.34 |
| Asp | moderate | minus | 17 | 4029.30 | 338.36 |
| Asp | severe | minus | 18 | 3932.58 | 279.15 |
| Asp | well-watered | plus | 18 | 3555.51 | 212.32 |
| Asp | moderate | plus | 19 | 3188.04 | 225.62 |
| Asp | severe | plus | 16 | 3457.92 | 370.01 |
| Gln | well-watered | minus | 18 | 2792.03 | 377.93 |
| Gln | moderate | minus | 17 | 2289.50 | 527.63 |
| Gln | severe | minus | 18 | 3963.96 | 737.07 |
| Gln | well-watered | plus | 18 | 2977.45 | 285.87 |

| | | | | | |
|---|---|---|---|---|---|
| Gln | moderate | plus | 19 | 2574.78 | 259.06 |
| Gln | severe | plus | 16 | 4116.11 | 772.69 |
| Glu | well-watered | minus | 18 | 18642.98 | 851.60 |
| Glu | moderate | minus | 17 | 15496.47 | 940.69 |
| Glu | severe | minus | 18 | 17833.28 | 1101.27 |
| Glu | well-watered | plus | 18 | 19349.94 | 724.37 |
| Glu | moderate | plus | 19 | 15347.24 | 797.09 |
| Glu | severe | plus | 16 | 17040.75 | 841.88 |
| His | well-watered | minus | 18 | 171.99 | 18.66 |
| His | moderate | minus | 17 | 113.73 | 9.20 |
| His | severe | minus | 18 | 308.10 | 61.75 |
| His | well-watered | plus | 18 | 243.88 | 25.33 |
| His | moderate | plus | 19 | 187.63 | 26.15 |
| His | severe | plus | 16 | 295.11 | 43.48 |
| Ile | well-watered | minus | 18 | 557.64 | 84.94 |
| Ile | moderate | minus | 17 | 235.60 | 32.92 |
| Ile | severe | minus | 18 | 784.52 | 206.07 |
| Ile | well-watered | plus | 18 | 802.64 | 110.58 |
| Ile | moderate | plus | 19 | 410.49 | 67.75 |
| Ile | severe | plus | 16 | 697.08 | 130.37 |
| Leu | well-watered | minus | 18 | 421.82 | 86.41 |
| Leu | moderate | minus | 17 | 213.50 | 36.73 |
| Leu | severe | minus | 18 | 1035.02 | 361.36 |
| Leu | well-watered | plus | 18 | 706.91 | 109.06 |
| Leu | moderate | plus | 19 | 526.00 | 140.93 |
| Leu | severe | plus | 16 | 946.00 | 186.18 |
| Lys | well-watered | minus | 18 | 70.28 | 8.03 |
| Lys | moderate | minus | 17 | 33.93 | 2.82 |
| Lys | severe | minus | 18 | 76.42 | 19.62 |
| Lys | well-watered | plus | 18 | 105.98 | 14.19 |
| Lys | moderate | plus | 19 | 46.03 | 3.54 |

| | | | | | |
|---|---|---|---|---|---|
| Lys | severe | plus | 16 | 85.45 | 19.55 |
| Met | well-watered | minus | 18 | 19.80 | 1.85 |
| Met | moderate | minus | 17 | 9.84 | 0.65 |
| Met | severe | minus | 18 | 13.34 | 2.28 |
| Met | well-watered | plus | 18 | 24.80 | 2.44 |
| Met | moderate | plus | 19 | 11.99 | 0.71 |
| Met | severe | plus | 16 | 17.35 | 1.80 |
| Phe | well-watered | minus | 18 | 452.15 | 67.73 |
| Phe | moderate | minus | 17 | 213.16 | 26.17 |
| Phe | severe | minus | 18 | 384.97 | 112.75 |
| Phe | well-watered | plus | 18 | 433.19 | 59.07 |
| Phe | moderate | plus | 19 | 211.78 | 24.16 |
| Phe | severe | plus | 16 | 300.15 | 42.43 |
| Pro | well-watered | minus | 18 | 462.38 | 56.06 |
| Pro | moderate | minus | 17 | 216.33 | 18.98 |
| Pro | severe | minus | 18 | 381.67 | 85.75 |
| Pro | well-watered | plus | 18 | 824.53 | 98.66 |
| Pro | moderate | plus | 19 | 525.54 | 53.43 |
| Pro | severe | plus | 16 | 752.64 | 97.41 |
| Ser | well-watered | minus | 18 | 5880.99 | 741.57 |
| Ser | moderate | minus | 17 | 2669.41 | 288.98 |
| Ser | severe | minus | 18 | 2445.13 | 222.82 |
| Ser | well-watered | plus | 18 | 3998.54 | 510.17 |
| Ser | moderate | plus | 19 | 1434.37 | 172.71 |
| Ser | severe | plus | 16 | 1632.69 | 161.49 |
| Thr | well-watered | minus | 18 | 1388.56 | 87.42 |
| Thr | moderate | minus | 17 | 1100.64 | 71.55 |
| Thr | severe | minus | 18 | 1303.13 | 155.85 |
| Thr | well-watered | plus | 18 | 1406.66 | 123.67 |
| Thr | moderate | plus | 19 | 873.72 | 88.38 |
| Thr | severe | plus | 16 | 1151.31 | 118.27 |

| | | | | | |
|---|---|---|---|---|---|
| Trp | well-watered | minus | 18 | 260.30 | 43.07 |
| Trp | moderate | minus | 17 | 213.83 | 36.86 |
| Trp | severe | minus | 18 | 566.29 | 118.33 |
| Trp | well-watered | plus | 18 | 614.56 | 65.35 |
| Trp | moderate | plus | 19 | 690.45 | 84.45 |
| Trp | severe | plus | 16 | 832.91 | 114.17 |
| Tyr | well-watered | minus | 18 | 153.68 | 15.07 |
| Tyr | moderate | minus | 17 | 91.04 | 8.34 |
| Tyr | severe | minus | 18 | 137.13 | 22.58 |
| Tyr | well-watered | plus | 18 | 199.06 | 17.70 |
| Tyr | moderate | plus | 19 | 127.47 | 10.40 |
| Tyr | severe | plus | 16 | 234.76 | 50.64 |
| Val | well-watered | minus | 18 | 881.07 | 123.50 |
| Val | moderate | minus | 17 | 365.43 | 44.30 |
| Val | severe | minus | 18 | 1002.19 | 259.77 |
| Val | well-watered | plus | 18 | 1218.31 | 180.53 |
| Val | moderate | plus | 19 | 600.13 | 93.31 |
| Val | severe | plus | 16 | 1022.28 | 187.39 |

**Abbreviations:** Ala, alanine; Asp, aspartic acid; Asn, asparagine; Glu, glutamic acid; Gln, glutamine; His, histidine; Ile, isoleucine; Leu, leucine; Met, methionine; Phe, phenylalanine; Pro, proline; Ser, serine; Thr, threonine; Trp, tryptophan; Tyr, tyrosine; Val, valine.

**Table S8. Statistical results for linear mixed-effects models testing the effects of water regime, herbivory, and their interaction on free amino acid concentrations.** Linear mixed-effects models included water regime, herbivory, and their interaction as fixed effects, and genotype as a random effect. Type III ANOVA was applied. Degrees of freedom were estimated using the Satterthwaite approximation. The table reports F-values and p-values, marginal and conditional coefficients of determination ($R^2m/R^2c$), and the transformations applied to each response variable. Significant effects are shown in bold ($p < 0.05$). Sample size: n = 16-18.

| | | **Water regime** | | **Herbivory** | | **Interaction** | | |
|---|---|---|---|---|---|---|---|---|
| **Response** | **Transformation** | **F-statistic** | **p-value** | **F-statistic** | **p-value** | **F-statistic** | **p-value** | **$R^2m/R^2c$** |
| Ala | sqrt | 4.12 | **0.019** | 1.60 | 0.209 | 1.23 | 0.296 | 0.11 / 0.13 |
| Arg | log | 43.66 | **0.000** | 3.20 | 0.076 | 3.30 | **0.041** | 0.38 / 0.60 |
| Asn | log | 18.57 | **0.000** | 3.80 | 0.054 | 0.55 | 0.577 | 0.22 / 0.44 |
| Asp | none | 0.17 | 0.841 | 6.52 | **0.012** | 0.72 | 0.488 | 0.06 / 0.22 |
| Gln | log | 4.17 | 0.018 | 2.41 | 0.123 | 0.18 | 0.839 | 0.09 / NA |
| Glu | none | 9.65 | 0.000 | 0.01 | 0.909 | 0.40 | 0.672 | 0.16 / 0.19 |
| His | log | 13.84 | **0.000** | 9.97 | **0.002** | 1.75 | 0.178 | 0.26 / 0.32 |
| Ile | log | 11.91 | **0.000** | 7.83 | **0.006** | 1.37 | 0.259 | 0.23 / 0.27 |
| Leu | log | 10.89 | 0.000 | 14.73 | **0.000** | 1.72 | 0.183 | 0.25 / 0.33 |
| Lys | log | 14.93 | 0.000 | 7.40 | **0.008** | 0.19 | 0.825 | 0.26 / NA |
| Met | log | 35.09 | **0.000** | 14.88 | **0.000** | 0.24 | 0.791 | 0.42 / 0.49 |
| Phe | log | 16.51 | 0.000 | 0.01 | 0.909 | 0.08 | 0.924 | 0.23 / 0.28 |
| Pro | sqrt | 13.84 | **0.000** | 61.04 | **0.000** | 0.13 | 0.881 | 0.39 / 0.53 |
| Ser | log | 51.80 | 0.000 | 34.99 | **0.000** | 0.74 | 0.477 | 0.49 / 0.64 |
| Thr | sqrt | 10.61 | 0.000 | 3.19 | **0.077** | 0.91 | 0.406 | 0.18 / 0.30 |
| Trp | sqrt | 7.07 | **0.001** | 52.19 | **0.000** | 1.83 | 0.165 | 0.38 / 0.42 |
| Tyr | log | 8.94 | 0.000 | 16.96 | **0.000** | 0.47 | 0.627 | 0.25 / NA |
| Val | log | 13.14 | **0.000** | 7.11 | **0.009** | 0.49 | 0.612 | 0.23 / 0.27 |

**Abbreviations:** Ala, alanine; Asp, aspartic acid; Asn, asparagine; Glu, glutamic acid; Gln, glutamine; His, histidine; Ile, isoleucine; Leu, leucine; Met, methionine; Phe, phenylalanine; Pro, proline; Ser, serine; Thr, threonine; Trp, tryptophan; Tyr, tyrosine; Val, valine.

**Table S9. Overview of all identified volatile compounds.** A total of 86 VOCs were identified, belonging to seven major chemical classes. Volatiles were collected using a push-pull system with compressed air filtered through a charcoal filter entering a section of the plant enclosed in a roasting bag and withdrawn at the top through a volatile collection trap (VCT) containing Poropak-Q as an adsorbent. VCTs were eluted after sampling with 200 µl dichloromethane containing nonyl acetate as an internal standard (10 ng µl⁻¹). VOCs were analysed using GC-MS and GC-FID for compound identification and quantification. The table lists compound names, class, SMILES (Simplified Molecular Input Line Entry System), InChIKey (International Chemical Identifier Key) notation, retention times, match and reverse match values, and match probabilities (Prob.) according to NIST database.

| Compound | Class | SMILES | InChIKey | RT | Match | R.match | Prob. (%) |
|---|---|---|---|---|---|---|---|
| Benzaldehyde | Aromatics | C1=CC=C(C=C1)C=O | HUMNYLRZRPPJDN-UHFFFAOYSA-N | 8.03 | 786 | 796 | 68 |
| Benzylalcohol | Aromatics | C1=CC=C(C=C1)CO | WVDDGKGOMKODPV-UHFFFAOYSA-N | 9.85 | 851 | 916 | 38.5 |
| Salicylaldehyde | Aromatics | C1=CC=C(C(=C1)C=O)O | SMQUZDBALVYZAC-UHFFFAOYSA-N | 10.05 | 923 | 926 | 71.8 |
| methyl salicylic acid | Aromatics | COC(=O)C1=CC=CC=C1O | OSWPMRLSEDHDFF-UHFFFAOYSA-N | 13.7 | 916 | 929 | 71 |
| Eugenol | Aromatics | COC1=C(C=CC(=C1)CC=C)O | RRAFCDWBNXTKKO-UHFFFAOYSA-N | 17.33 | 827 | 883 | 40.4 |
| Jasmone | Aromatics | CC/C=C\CC1=C(CCC1=O)C | XMLSXPIVAXONDL-PLNGDYQASA-N | 18.27 | 822 | 829 | 74.8 |
| Benzoic acid, 2-methylbutyl ester | Aromatics | CCC(C)COC(=O)C1=CC=CC=C1 | PYZHESNNAPENLQ-UHFFFAOYSA-N | 19.01 | 907 | 916 | 53.7 |
| 3-Hexenal | GLV | CC/C=C/CC=O | GXANMBISFKBPEX-ONEGZZNKSA-N | 4.63 | 884 | 931 | 41.8 |
| 3-Hexenol | GLV | CC/C=C\\CCO | UFLHIIWVXFIJGU-ARJAWSKDSA-N | 5.75 | 932 | 943 | 79.8 |
| 3-Hexen-1-ol, acetate | GLV | CC/C=C/CCOC(=O)C | NPFVOOAXDOBMCE-SNAWJCMRSA-N | 9.2 | 939 | 963 | 51.8 |
| 3-Hexen-1-ol, acetate_2 | GLV | CC/C=C/CCOC(=O)C | NPFVOOAXDOBMCE-SNAWJCMRSA-N | 9.37 | 752 | 829 | 20.2 |
| trans-3-Hexenyl butyrate | GLV | CCCC(=O)OCC/C=C/CC | ZCHOPXVYTWUHDS-AATRIKPKSA-N | 13.48 | 807 | 909 | 35.3 |
| cis-3-hexenyl-α-methylbutyrate_1 | GLV | CC/C=C\\CCOC(=O)C(C)CC | JKKGTSUICJWEKB-SREVYHEPSA-N | 14.56 | 822 | 875 | 31.5 |
| cis-hexenyl isovalerate | GLV | CC/C=C\CCOC(=O)CC(C)C | AIQLNKITFBJPFO-WAYWQWQTSA-N | 14.64 | 814 | 892 | 49.9 |
| cis-hexenyl valerate | GLV | CCCCC(=O)OCC/C=C\CC | XPFTVTFOOTVHIA-ALCCZGGFSA-N | 14.88 | 614 | 743 | 12.6 |
| cis-3-hexenyl-α-methylbutyrate_2 | GLV | CC/C=C\\CCOC(=O)C(C)CC | JKKGTSUICJWEKB-SREVYHEPSA-N | 14.97 | 628 | 767 | 45.4 |
| cis-3-hexenylbenzoate | GLV | CC/C=C\\CCOC(=O)C1=CC=CC=C1 | BCOXBEHFBZOJJZ-ARJAWSKDSA-N | 21.65 | 869 | 917 | 65.8 |
| unknown_GLV_1 | GLV | NA | NA | 21.77 | 590 | 590 | 45.1 |
| DMNT/Phenylethylalcohol | Homoterpene | CC(=CCCC(=CC=C)C)C | LUKZREJJLWEWQM-UHFFFAOYSA-N | 11.82 | 900 | 915 | 80 |
| α-Pinene | Monoterpenes | CC1=CCC2CC1C2(C)C | GRWFGVWFFZKLTI-UHFFFAOYSA-N | 7.38 | 881 | 930 | 14.8 |
| Camphene | Monoterpenes | CC1(C2CCC(C2)C1=C)C | CRPUJAZIXJMDBK-UHFFFAOYSA-N | 7.73 | 879 | 927 | 33.1 |
| Sabinene | Monoterpenes | CC(C)C12CCC(=C)C1C2 | NDVASEGYNIMXJL-UHFFFAOYSA-N | 8.34 | 901 | 914 | 24.9 |
| β-Pinene | Monoterpenes | CC1(C2CCC(=C)C1C2)C | WTARULDDTDQWMU-UHFFFAOYSA-N | 8.4 | 854 | 913 | 20.8 |
| β-Myrcene | Monoterpenes | CC(=CCCC(=C)C=C)C | UAHWPYUMFXYFJY-UHFFFAOYSA-N | 8.76 | 703 | 833 | 38.4 |

| | | | | | | | |
|---|---|---|---|---|---|---|---|
| Limonene | Monoterpenes | CC1=CCC(CC1)C(=C)C | XMGQYMWWDOXHJM-UHFFFAOYSA-N | 9.68 | 746 | 842 | 31.6 |
| 1,8 Cineol | Monoterpenes | CC1(C2CCC(O1)(CC2)C)C | WEEGYLXZBRQIMU-UHFFFAOYSA-N | 9.8 | 913 | 927 | 83.2 |
| trans-β-Ocimene | Monoterpenes | CC(=CC/C=C(\\C)/C=C)C | IHPKGUQCSIINRJ-CSKARUKUSA-N | 9.91 | 959 | 959 | 24.2 |
| (*E*)-β-Ocimene | Monoterpenes | CC(=CCC=C(C)C=C)C | IHPKGUQCSIINRJ-UHFFFAOYSA-N | 10.16 | 955 | 955 | 24.2 |
| Linalool | Monoterpenes | CC(=CCCC(C)(C=C)O)C | CDOSHBSSFJOMGT-UHFFFAOYSA-N | 11.44 | 933 | 933 | 70.7 |
| α-Terpineol | Monoterpenes | CC1=CCC(CC1)C(C)(C)O | WUOACPNHFRMFPN-UHFFFAOYSA-N | 13.64 | 831 | 831 | 27.4 |
| β-Cyclocitral | Monoterpenes | CC1=C(C(CCC1)(C)C)C=O | MOQGCGNUWBPGTQ-UHFFFAOYSA-N | 14.33 | 790 | 818 | 68 |
| Butyl aldoxime, 3-methyl-, syn- | N | CC(C)C/C=N\\O | JAUPRNSQRRCCRR-XQRVVYSFSA-N | 5.6 | 766 | 821 | 39.8 |
| Butyl aldoxime, 2-methyl-, anti- | N | CCC(C)C=NO | SEWWFHKIKWFJNV-UHFFFAOYSA-N | 5.91 | 900 | 909 | 50.9 |
| Butyl aldoxime, 2-methyl-, syn- | N | CCC(C)/C=N\\O | SEWWFHKIKWFJNV-XQRVVYSFSA-N | 6 | 917 | 926 | 80.1 |
| Butyl aldoxime, 3-methyl-, anti-_2 | N | CC(C)C/C=N\\O | JAUPRNSQRRCCRR-XQRVVYSFSA-N | 6.07 | 898 | 909 | 70.4 |
| Nitropentane | N | CCCCC[N+](=O)[O-] | BVALZCVRLDMXOQ-UHFFFAOYSA-N | 6.69 | 784 | 885 | 39.1 |
| Benzyl cyanide | N | C1=CC=C(C=C1)CC#N | SUSQOBVLVYHIEX-UHFFFAOYSA-N | 12.35 | 961 | 967 | 47.8 |
| o-Tolyl isocyanide | N | CC1=CC=CC=C1[N+]#[C-] | HGHZICGHCZFYNX-UHFFFAOYSA-N | 12.58 | 814 | 877 | 27.8 |
| Benzyl isocyanide | N | [C-]#[N+]CC1=CC=CC=C1 | RIWNFZUWWRVGEU-UHFFFAOYSA-N | 12.7 | 799 | 855 | 26 |
| unknown_N_1 | N | NA | NA | 13 | 499 | 855 | 26 |
| unknown_N_3 | N | NA | NA | 15.03 | 576 | 711 | 25.5 |
| 4-Methylbenzaldoxime_1 | N | CC1=CC=C(C=C1)/C=N/O | SRNDYVBEUZSFEZ-RMKNXTFCSA-N | 15.15 | 788 | 882 | 81.8 |
| 4-Methylbenzaldoxime_2 | N | CC1=CC=C(C=C1)/C=N/O | SRNDYVBEUZSFEZ-RMKNXTFCSA-N | 15.47 | 712 | 849 | 44.1 |
| unknown_N_2 | N | NA | NA | 15.55 | 549 | 621 | 17.8 |
| Indol | N | C1=CC=C2C(=C1)C=CN2 | SIKJAQJRHWYJAI-UHFFFAOYSA-N | 15.94 | 906 | 908 | 50.5 |
| 2-nitroethyl benzene | N | C1=CC=C(C=C1)CC[N+](=O)[O-] | XAWCLWKTUKMCMO-UHFFFAOYSA-N | 16.03 | 886 | 922 | 43 |
| 4-ethyl-1-hexene | other | CCC(CC)CC=C | OPMUAJRVOWSBTP-UHFFFAOYSA-N | 6.18 | 878 | 884 | 58.4 |
| 4-Penten-1-ol acetate | other | CC(=O)OCCCC=C | LVHDNIMNOMRZMF-UHFFFAOYSA-N | 6.35 | 848 | 928 | 71.8 |
| 2-ethyl Butanol | other | CCC(CC)CO | TZYRSLHNPKPEFV-UHFFFAOYSA-N | 6.61 | 786 | 892 | 28.3 |
| 5-Hepten-2-one, 6-methyl | other | CC(=CCCC(=O)C)C | UHEPJGULSIKKTP-UHFFFAOYSA-N | 8.58 | 876 | 924 | 23.1 |
| 2-Hydroxy-Cyclohexanone | other | C1CCC(=O)C(C1)O | ODZTXUXIYGJLMC-UHFFFAOYSA-N | 8.91 | 858 | 897 | 82.9 |
| 2-Cyclohexenone-4-hydroxy | other | C1CC(=O)C=CC1O | AMFCFGFZMSQOIU-UHFFFAOYSA-N | 9.04 | 698 | 773 | 33 |
| Tetrahydrocyclopenta[1,3]dioxin-4-one | other | C1CC2C(C1)OCOC2=O | UNLDNDXCXJPPOK-UHFFFAOYSA-N | 9.44 | 748 | 756 | 46.4 |
| Tetrahydrocyclopenta[1,3]dioxin-4-one_2 | other | C1CC2C(C1)OCOC2=O | UNLDNDXCXJPPOK-UHFFFAOYSA-N | 9.51 | 743 | 760 | 43.5 |

| | | | | | | | |
|---|---|---|---|---|---|---|---|
| 2,6-Cyclooctadien-1-ol | other | C/1C/C=C\\C(C/C=C1)O | FRIHCVHTOVWRBC-ILNZVKDWSA-N | 10.65 | 798 | 826 | 50.2 |
| α-Methyl-α-(4-methyl-3-pentenyl)-oxiranemethanol | other | CC(=CCCC(C)(C1CO1)O)C | BXOURKNXQXLKRK-UHFFFAOYSA-N | 10.83 | 868 | 888 | 30.4 |
| unknown_other_1 | other | NA | NA | 10.96 | 513 | 702 | 25.8 |
| Linalool oxide | other | CC1(CCC(O1)C(C)(C)O)C=C | BRHDDEIRQPDPMG-UHFFFAOYSA-N | 11.21 | 703 | 713 | 20.2 |
| Nonanal | other | CCCCCCCCC=O | GYHFUZHODSMOHU-UHFFFAOYSA-N | 11.54 | 877 | 893 | 56.3 |
| Borneol | other | CC1(C2CCC1(C(C2)O)C)C | DTGKSKDOIYIVQL-UHFFFAOYSA-N | 13.13 | 857 | 877 | 56.3 |
| Decanal | other | CCCCCCCCCC=O | KSMVZQYAVGTKIV-UHFFFAOYSA-N | 13.94 | 903 | 908 | 68.1 |
| Heptylcyclohexane | other | CCCCCCCC1CCCCC1 | MSTLSCNJAHAQNU-UHFFFAOYSA-N | 16.61 | 790 | 826 | 15.1 |
| α-Cubebene | Sesquiterpenes | C[C@@H]1CC[C@H]([C@H]2[C@]13[C@@H]2C(=CC3)C)C(C)C | XUEHVOLRMXNRKQ-KHMAMNHCSA-N | 17.18 | 871 | 888 | 26.5 |
| Copaene_1 | Sesquiterpenes | CC1=CC[C@H]2[C@H]3[C@@H]1[C@@]2(CC[C@H]3C(C)C)C | VLXDPFLIRFYIME-BTFPBAQTSA-N | 17.76 | 881 | 893 | 26.5 |
| β-Cubebene | Sesquiterpenes | C[C@@H]1CC[C@H]([C@H]2[C@]13[C@@H]2C(=C)CC3)C(C)C | FSRZGYRCMPZNJF-KHMAMNHCSA-N | 18.05 | 654 | 751 | 28.1 |
| unknown_sesquiterpene_1 | Sesquiterpenes | NA | NA | 18.09 | 930 | 932 | 25.7 |
| (*E*)-β-Caryophyllene | Sesquiterpenes | C/C/1=C\\CCC(=C)[C@@H]2CC([C@H]2CC1)(C)C | NPNUFJAVOOONJE-IOMPXFEGSA-N | 18.69 | 943 | 945 | 37.5 |
| β-Copaene | Sesquiterpenes | CC(C)[C@@H]1CC[C@]2([C@@H]3[C@H]1C2C(=C)CC3)C | UPVZPMJSRSWJHQ-XIQJJJERSA-N | 18.87 | 787 | 804 | 15.3 |
| cis-Muurola-4(15),5-diene_1 | Sesquiterpenes | C[C@@H]1CC[C@@H](C2=CC(=C)CCC12)C(C)C | RNDFUOKDULDZPR-ZFXTZCCVSA-N | 19.3 | 891 | 933 | 29.7 |
| α-Humulene | Sesquiterpenes | C/C/1=C\\CC(/C=C/C/C(=C/CC1)/C)(C)C | FAMPSKZZVDUYOS-HRGUGZIWSA-N | 19.39 | 827 | 840 | 54.9 |
| Alloaromadendrene | Sesquiterpenes | C[C@@H]1CC[C@H]2[C@@H]1[C@H]3[C@H](C3(C)C)CCC2=C | ITYNGVSTWVVPIC-DHGKCCLASA-N | 19.54 | 884 | 899 | 16.9 |
| cis-Muurola-4(15),5-diene_2 | Sesquiterpenes | C[C@@H]1CC[C@@H](C2=CC(=C)CCC12)C(C)C | RNDFUOKDULDZPR-ZFXTZCCVSA-N | 19.58 | 657 | 757 | 48.6 |
| y-Muurolene | Sesquiterpenes | CC1=C[C@@H]2[C@H](CC1)C(=C)CC[C@H]2C(C)C | WRHGORWNJGOVQY-ZNMIVQPWSA-N | 19.83 | 880 | 891 | 22.5 |
| Germacrene D | Sesquiterpenes | C/C/1=C\\CCC(=C)/C=C/[C@@H](CC1)C(C)C | GAIBLDCXCZKKJE-RXJOXMPGSA-N | 19.94 | 896 | 918 | 33.9 |
| unknown_sesquiterpene_2 | Sesquiterpenes | NA | NA | 20.05 | 896 | 903 | 20.1 |
| unknown_sesquiterpene_3 | Sesquiterpenes | NA | NA | 20.16 | 837 | 847 | 7.57 |
| Copaene_2 | Sesquiterpenes | CC1=CC[C@H]2[C@H]3[C@@H]1[C@@]2(CC[C@H]3C(C)C)C | VLXDPFLIRFYIME-BTFPBAQTSA-N | 20.22 | 774 | 821 | 9.51 |
| α-Muurolene | Sesquiterpenes | CC1=C[C@@H]2[C@H](CC1)C(=CC[C@H]2C(C)C)C | QMAYBMKBYCGXDH-ZNMIVQPWSA-N | 20.29 | 906 | 909 | 21.1 |
| α-Farnesene | Sesquiterpenes | CC(=CCC/C(=C/C/C=C(\\C)/C=C)/C)C | CXENHBSYCFFKJS-VDQVFBMKSA-N | 20.4 | 952 | 960 | 65 |
| γ-Cadinene | Sesquiterpenes | CC1=C[C@@H]2[C@@H](CC1)C(=C)CC[C@H]2C(C)C | WRHGORWNJGOVQY-KKUMJFAQSA-N | 20.58 | 927 | 931 | 34.5 |
| δ-Cadinene | Sesquiterpenes | CC1=C[C@H]2[C@@H](CCC(=C2CC1)C)C(C)C | FUCYIEXQVQJBKY-ZFWWWQNUSA-N | 20.75 | 908 | 908 | 60.8 |
| Cubenene | Sesquiterpenes | C12C3C4C1=C5C2C3=C45 | KWFAQPWLROZBAY-UHFFFAOYSA-N | 20.93 | 765 | 797 | 28.3 |
| α-Cadinene | Sesquiterpenes | CC1=C[C@@H]2[C@@H](CC1)C(=CC[C@H]2C(C)C)C | QMAYBMKBYCGXDH-KKUMJFAQSA-N | 21.03 | 844 | 870 | 27.3 |
| unknown_sesquiterpene_4 | Sesquiterpenes | NA | NA | 21.49 | 841 | 854 | 26.9 |

| | | | | | | | |
|---|---|---|---|---|---|---|---|
| Caryophyllene oxide | Sesquiterpenes | C[C@@]12CC[C@@H]3[C@H](CC3(C)C)C(=C)CC[C@H]1O2 | NVEQFIOZRFFVFW-RGCMKSIDSA-N | 21.96 | 872 | 873 | 55.8 |
| τCadinol | Sesquiterpenes | CC1=C[C@H]2[C@@H](CC[C@]([C@@H]2CC1)(C)O)C(C)C | LHYHMMRYTDARSZ-XQLPTFJDSA-N | 23 | 885 | 887 | 47.3 |

**Table S10. Sample sizes and number of detected volatile organic compounds (VOCs) across treatments.** Shown are water regime (field capacity, %), herbivory treatment, number of samples analysed (n_samples), and number of detected VOCs (n_compounds_detected).

| Water regime | Herbivory | n_samples | n_compounds_detected |
|---|---|---|---|
| 100 | minus | 18 | 83 |
| 100 | plus | 18 | 86 |
| 50 | minus | 18 | 81 |
| 50 | plus | 18 | 85 |
| 25 | minus | 18 | 74 |
| 25 | plus | 16 | 83 |

**Table S11. Statistical results for linear mixed-effects models testing the effects of water regime, herbivory, and their interaction on total VOC emissions, VOC classes, and functional Hill diversity ($q_1$).** Linear mixed-effects models included water regime, herbivory, and their interaction as fixed effects, and genotype as a random effect. Type III ANOVA was applied. Degrees of freedom were estimated using the Satterthwaite approximation. The table reports F-values and p-values, marginal and conditional coefficients of determination ($R^2m/R^2c$), and the transformations applied to each response variable. Significant effects are shown in bold ($p < 0.05$). Sample size: n = 16-18.

| | | Water regime | | Herbivory | | Interaction | | |
|---|---|---|---|---|---|---|---|---|
| **Response** | **Transformation** | **F-statistic** | **p-value** | **F-statistic** | **p-value** | **F-statistic** | **p-value** | **$R^2m/R^2c$** |
| **VOCs** | | | | | | | | |
| Total emission | log | 0.97 | 0.384 | 99.16 | **0.000** | 0.33 | 0.722 | 0.48 / 0.51 |
| Aromatics | log | 0.95 | 0.390 | 176.77 | **0.000** | 1.13 | 0.326 | 0.63 / NA |
| GLVs | log | 0.42 | 0.658 | 83.02 | **0.000** | 0.11 | 0.899 | 0.43 / 0.46 |
| DMNT | log | 3.20 | **0.045** | 127.15 | **0.000** | 2.94 | 0.057 | 0.56 / 0.59 |
| Monoterpenes | log | 2.94 | 0.057 | 77.15 | **0.000** | 1.33 | 0.270 | 0.44 / 0.46 |
| Sesquiterpenes | log | 0.98 | 0.379 | 22.15 | **0.000** | 0.04 | 0.962 | 0.18 / 0.25 |
| Nitrogeneous compounds | log | 2.13 | 0.124 | 165.98 | **0.000** | 0.40 | 0.674 | 0.61 / 0.62 |
| Others | log | 2.18 | 0.118 | 129.91 | **0.000** | 3.59 | **0.031** | 0.57 / 0.58 |
| **Functional Hill diversity** | | | | | | | | |
| Functional $q_1$ Shannon | | | | | | 6.45 | **0.002** | |

**Table S12. Statistical results for linear mixed-effects models testing the effects of water regime, herbivory, and their interaction on selected volatile organic compounds.** Linear mixed-effects models included water regime, herbivory, and their interaction as fixed effects, and genotype as a random effect. Type III ANOVA was applied with Satterthwaite approximation of degrees of freedom. The table reports F-values, FDR-adjusted p-values, marginal and conditional coefficients of determination ($R^2m/R^2c$), and the transformations applied to each response variable. Significant effects are shown in bold ($p < 0.05$). Sample size: n = 16-18.

| | | **Water regime** | | **Herbivory** | | **Interaction** | | |
|---|---|---|---|---|---|---|---|---|
| **Response** | **Transformation** | **F-statistic** | **p-value** | **F-statistic** | **p-value** | **F-statistic** | **p-value** | **$R^2m/R^2c$** |
| 3-Hexenal | log | 5.35 | **0.010** | 160.02 | **0.000** | 1.30 | 0.439 | 0.36 /0.37 |
| α-Pinene | log | 6.25 | **0.006** | 20.20 | **0.000** | 0.33 | 0.791 | 0.15 /0.16 |
| (*E*)-β-Ocimene | log | 6.45 | **0.006** | 323.58 | **0.000** | 2.04 | 0.242 | 0.51 /0.55 |
| Benzyl cyanide | log | 6.70 | **0.006** | 1.14 | 0.289 | 1.02 | 0.532 | 0.64 /0.65 |
| Benzaldehyde | log | 6.36 | **0.006** | 56.57 | **0.000** | 3.21 | 0.121 | 0.31 /0.33 |
| Salicylaldehyde | log | 0.16 | 0.855 | 180.07 | **0.000** | 2.82 | 0.152 | 0.76 / NA |

**Table S13. Statistical results for linear mixed-effects regression models testing the effects of leaf damage intensity, water regime, and their interaction on individual VOC emissions.** Linear mixed-effects regression models included leaf damage intensity (%), water regime, and their interaction as fixed effects, and genotype as a random effect. Type III ANOVA was applied. Degrees of freedom were estimated using the Satterthwaite approximation. The table reports FDR-adjusted p-values, and marginal and conditional coefficients of determination ($R^2m/R^2c$). Significant effects are shown in bold ($p < 0.05$). Sample size: n = 16-18.

| | | **Water regime** | | **Herbivory** | | **Interaction** | | |
|---|---|---|---|---|---|---|---|---|
| **Response** | **Transformation** | **F-statistic** | **p-value** | **F-statistic** | **p-value** | **F-statistic** | **p-value** | **$R^2m/R^2c$** |
| 3-Hexenal | log | 0.17 | 0.845 | 0.97 | 0.329 | 0.60 | 0.553 | 0.13 / NA |
| α-Pinene | log | 6.66 | **0.003** | 7.28 | **0.011** | 10.66 | **0.000** | 0.40 / 0.42 |
| (*E*)-β-Ocimene | log | 7.59 | **0.001** | 29.19 | **0.000** | 6.95 | **0.002** | 0.46 / NA |
| Benzyl cyanide | log | 6.03 | **0.004** | 12.91 | **0.001** | 3.92 | **0.026** | 0.35 / NA |
| Benzaldehyde | log | 6.38 | **0.003** | 12.37 | **0.001** | 6.37 | **0.003** | 0.34 / 0.44 |
| Salicylaldehyde | log | 2.86 | 0.066 | 35.46 | **0.000** | 7.19 | **0.002** | 0.47 / NA |
